# Rewiring central carbon metabolism reveals a trade-off between yield and production rate in poly-γ-glutamic acid synthesis by *Bacillus subtilis*

**DOI:** 10.64898/2026.10.19.756727

**Authors:** Frederik Völker, Michael Kohlstedt, Birthe Halmschlag, Johanna Müller, Janina Weinert, Christoph Wittmann, Lars M. Blank

**Author notes:** Correspondence: Prof. Dr. Lars M. Blank, Institute of Applied Microbiology-iAMB, Aachen Biology and Biotechnology ABBt, RWTH Aachen University, Worringer Weg 1, 52074 Aachen, Germany.

## Abstract

Improving carbon yield and process time is a key requirement for the industrially viable production of biopolymers. During poly-γ-glutamic acid production with *Bacillus subtilis*, a considerable fraction of the carbon source is diverted into by-products. These originate from accumulating pyruvate, constraining carbon flux towards glutamate synthesis. Disrupting by-product pathways reduces their secretion, but the underlying imbalance at the pyruvate node persists.

Here, we rewired the central carbon metabolism to redirect glycolytic flux from pyruvate formation towards oxaloacetate and 2-oxoglutarate in an improved medium. Deletion of pyruvate kinase combined with heterologous expression of phosphoenolpyruvate carboxylase established an alternative anaplerotic entry route into the TCA cycle. Subsequent disruption of gluconeogenetic reactions (OF5) reduced by-product formation by 81% and increased γ-PGA yield to 0.41 g g⁻¹ glucose, but lowered the glucose uptake rate. Alternatively, disruption of the TCA cycle downstream of 2-oxoglutarate combined with enhanced glutamate synthesis (OF4.2) increased substrate uptake and improved the γ-PGA production rate by 39% compared to the wild type, while product and by-product titers remained comparable. Both strategies thus revealed a trade-off between improved yield or production rate. Subsequent ¹³C metabolic flux analysis and intracellular metabolite profiling revealed distinct carbon routing phenotypes. While strain OF5 maintained carbon within the TCA cycle, OF4.2 cycled carbon between oxaloacetate, malate and pyruvate, thereby altering redox and energy balances. These results highlight the need for coordinated control of carbon flux, redox metabolism and energy supply for improving γ-PGA synthesis, and identify the oxaloacetate–malate–pyruvate node as a promising strain engineering target.

## 1. Introduction

As the global demand for chemicals continues to rise, the integration of novel biotechnological production routes plays a complementary and increasingly vital role alongside the chemical industry’s ongoing transition toward green and sustainable practices (Mesa et al., 2024; Sheldon and Brady, 2022). While traditional chemical synthesis has made significant advances in improving efficiency and reducing environmental impact, biotechnology offers additional options by enabling the use of renewable feedstocks, milder reaction conditions, and flexible systems that can be tailored for selective, low-emission production (Ioannidou et al., 2020). The development of microbial platforms capable of producing bulk and fine chemicals from sugars, CO₂, or industrial by-products aligns with the principles of green chemistry and supports the diversification and decarbonization of chemical supply chains towards a circular, bio-based economy (Ewing et al., 2022). This transformation is crucial not only for climate change mitigation, but also for long-term economic and resource resilience.

The Gram-positive bacterium *Bacillus subtilis*, one of the workhorses of biotechnology, plays a key role in this green transition. The fast-growing microbe has been subject of decades of research in molecular biology, genetics and physiology, resulting in a comprehensive understanding of its metabolism, regulatory systems, and stress response (Stülke et al., 2023). *B. subtilis* offers GRAS status and natively secrets proteins directly into the medium, simplifying downstream applications while also reducing energy demand and overall process costs (Chen et al., 2023). This, in combination with a plethora of available synthetic biology tools and metabolic models, makes this microorganism a promising chassis for sustainable bioengineering.

Bio-based polymers resemble key components of a future circular economy, given the urgent need for sustainable alternatives to currently used polymer materials of petrochemical origin. Among them, poly-γ-glutamic acid (γ-PGA) has gained significant attention as a candidate biomaterial due to its high biodegradability, non-toxicity and adjustable functional properties (Nair et al., 2023; Park et al., 2021). As a natural product of a variety of *Bacillus* strains, the polyamino acid is composed of different compositions of D- and L-glutamate linked via γ-peptide bonds, distinguishing the polymer from conventional α-peptides. The composition and genetic origin of the γ-PGA synthetase complex (PgsBCA) influence both the molecular weight and the ratio of D- and L-glutamate in the polymer, enabling customization of γ-PGA for applications ranging from medical to agricultural use (Halmschlag et al., 2019). For example, recent studies have presented γ-PGA as candidate for hydrogel formation and drug release as well as soil conditioner to improve water retention in sandy soils, highlighting the diversity of potential use cases (Balogun-Agbaje et al., 2021; Fan et al., 2017; Skalski et al., 2024).

One of the major challenges in the biotechnological production of γ-PGA is achieving a resource efficient production processes. Depending on the production strategy, γ-PGA synthesis can either rely on an external glutamate feed (glutamate-dependent processes) or on de novo glutamate synthesis from a carbon sources such as glucose (glutamate-independent processes). While glutamate-dependent processes can achieve comparably high γ-PGA titers of up to 75 g L^-1^, their reliance on external glutamate sources increases substrate costs (Jiang et al., 2016). In contrast, glutamate-independent processes are more economically favorable and can yield titers of up to 30 g L^-1^, but often feature an inherently low carbon yield and suboptimal production rate, especially under high-glucose fermentation conditions (Halmschlag et al., 2023; Peng et al., 2015). In many microbial hosts, excessive glucose availability can trigger overflow metabolism, a phenomenon where central carbon flux exceeds the capacity of the tricarboxylic acid (TCA) cycle and respiration (Li et al., 2022; Shimizu and Matsuoka, 2019). In this scenario, microbes incompletely oxidize their growth substrate (e.g., glucose) and secrete overflow metabolites or by-products, which, in case of *B. subtilis*, mainly consist of acetoin, 2,3-butanediol, acetate, and lactate (Völker et al., 2025). Although partial oxidation yields less ATP than full oxidation through respiration, overflow metabolism is a common metabolic strategy among fast growing cells. One possible explanation for the evolution of this mechanism across diverse organisms is microbial competition in natural environments, where rapid growth and efficient carbon source utilization provide a competitive advantage. In *E. coli*, it has been proposed that enzymes required for respiration are produced at higher costs compared to enzymes involved in partial oxidation of the carbon source, ultimately yielding a trade-off between resource allocation and fast growth (Basan et al., 2015; Molenaar et al., 2009). Notably, overflow metabolism is also required to maintain redox balance. Under conditions of high glycolytic flux, TCA cycle and respiration may become saturated, leading to an accumulation of NADH. By diverting excess pyruvate into overflow metabolites such as lactate and 2,3-butanediol, *B. subtilis* can effectively oxidize NADH to NAD^+^, allowing the microbe to adaptively balance its redox metabolism. While overflow metabolism offers a physiological advantage in natural environments, its presence remains undesirable in biotechnological processes. The secretion of partially oxidized by-products represents a significant carbon loss that could otherwise contribute to biomass or product synthesis, thereby lowering overall process efficiency. Moreover, the accumulation of organic acids leads to acidification of the culture medium, which can impair cell growth, inhibit enzyme activity, and trigger feedback inhibition of central metabolic pathways.

Multiple studies have already targeted reducing the overflow metabolism of *B. subtilis*. For example, the glyoxylate shunt of *B. licheniformis* was successfully introduced to *B. subtilis* to enable growth on overflow metabolites (Kabisch et al., 2013). Alternatively, Yuan et al. (2023) developed a quorum-sensing-based regulatory system to dynamically control glucose uptake, thereby mitigating the onset of overflow metabolism through demand-based substrate utilization. A widely adopted approach involves targeted deletion of genes encoding enzymes responsible for accumulation of overflow metabolites (Halmschlag et al., 2023). For example, Ma et al. (2018) inhibited acetoin synthesis by disrupting the relevant genetic pathway, which led to an increase in N-acetyl-D-glucosamine (GlcNAc) production. However, these strategies are not directly translatable to optimize γ-PGA production, as each optimization strategy depends on the specific biosynthetic demands of the product, such as redox demand, precursor availability, and energetic cost. Addressing this, we recently demonstrated a γ-PGA specific strategy by sequentially deleting genes involved in acetoin, acetate, lactate, and 2,3-butanediol formation in γ-PGA-producing *B. subtilis*, thereby reducing by-product accumulation while increasing product titer (Halmschlag et al., 2023). However, we observed that these pathway interruptions induced significant pyruvate accumulation and medium acidification, ultimately impairing the performance of the newly generated strains. These observations suggested that blocking individual by-product pathways without addressing the cause of overflow metabolism, namely the buildup of intracellular pyruvate, was insufficient. As most overflow metabolites are derived directly from pyruvate in *B. subtilis*, its accumulation serves as a metabolic bottleneck that triggers acid stress and redox imbalance.

The “push–pull–promote” concept introduced by Gu et al. (2019) was proven successful in avoiding the accumulation of overflow metabolites while increasing titer, yield and production rate of their product of interest GlcNAc. In their study, the authors successfully “pushed” carbon flux towards the GlcNAc synthesis pathway, “pulled” overflow carbon into the TCA cycle and “promoted” product synthesis by alleviating rate limiting steps and redox limitations.

Inspired by this framework, we adapted a similar systems-level design to modulate central carbon metabolism in favor of γ-PGA biosynthesis, with a particular focus on depleting excess pyruvate. First, we deleted the endogenous pyruvate kinase gene (*pykA*) to prevent the conversion of phosphoenolpyruvate (PEP) to pyruvate, thereby directly reducing pyruvate overaccumulation. To mitigate potential PEP accumulation and restore carbon flux into the tricarboxylic acid (TCA) cycle, a non-native phosphoenolpyruvate carboxylase (*ppc*) from *Corynebacterium glutamicum* was expressed heterologously, enabling the redirection of carbon flux from PEP to oxaloacetate (OAA). A similar study has been performed by Yokota et al. (2017) who investigated the effect of different *ppc* mutations on glutamic acid and lysine production in a pyruvate kinase deletion mutant. In this scenario and with glucose as carbon source, the phosphotransferase system still provides pyruvate from PEP to generate acetyl-CoA via pyruvate dehydrogenase (PDH), thereby ensuring TCA cycle functionality. In parallel, we reduced carbon loss by gluconeogenesis and futile cycling by disrupting reverse metabolic fluxes. In a subsequent step, we investigated whether redirecting flux toward glutamate synthesis could further enhance γ-PGA production by exploring a targeted disruption of the TCA cycle. Specifically, the operon of the 2-oxoglutarate dehydrogenase complex *odhAB* was replaced by a hyperactive variant of the transcriptional glutamate regulator *GltC*, thereby activating glutamate synthesis (via GltAB) and preventing further oxidation of 2-oxoglutarate in the TCA cycle. The respective engineering approaches were evaluated for their effect on glucose uptake, accumulation of overflow metabolites and production parameters of γ-PGA synthesis.

## 2. Materials and methods

### 2.1 Bacterial strains and growth conditions

All strains used in this study are listed in Table 1. Plasmids for *Bacillus* strain engineering were constructed in *Escherichia coli* DH5α, while the recombinase-positive strain *E. coli* JM101 was used to yield plasmids applicable for *B. subtilis* transformation. Plasmid construction, transformation and selection were performed as described in previous work (Völker et al., 2025). In brief, strains were regularly cultivated in LB medium at 37°C, containing (per liter): 10 g tryptone, 5 g yeast extract, and 10 g NaCl, adjusted to pH 7.4. If necessary, 100 µg mL⁻¹ spectinomycin or 0.5% (w/v) mannose was added to the medium. For solid media preparation, 20 g L⁻¹ agar was added. All complex media components were obtained from Carl Roth GmbH & Co. KG (Karlsruhe, Germany).

**Table 1:**
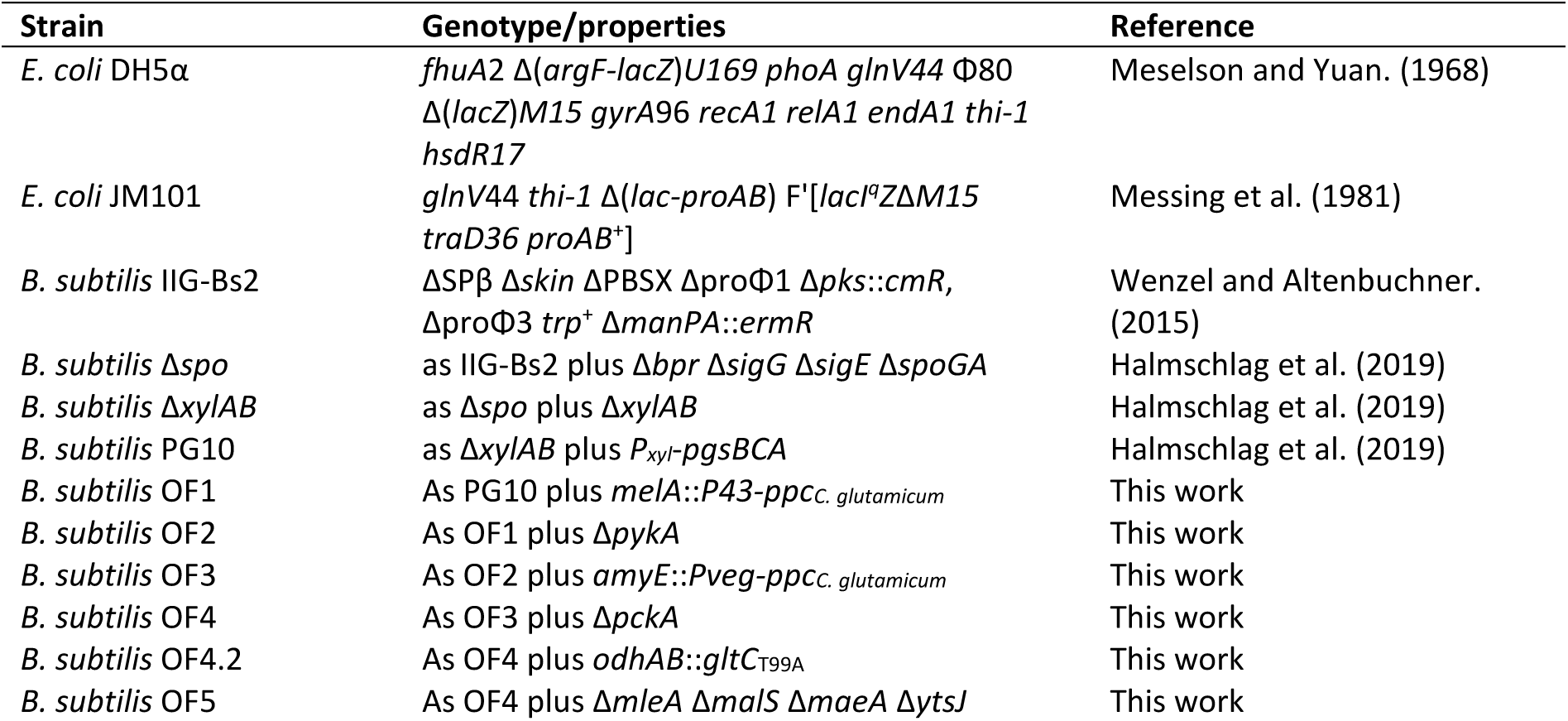
Strains used in this study.

### 2.2 Strain construction

Deletions and integrations were conducted as described previously. We used the sporulation-deficient strain *B. subtilis* Δ*spo*, which is derived from the *manPA*-negative strain *B. subtilis* IIG-Bs2, as starting point for genetic modifications (Halmschlag et al., 2020; Wenzel and Altenbuchner, 2015). Due to a frameshift mutation in the *swrA* allele, this strain is unable to produce γ-PGA natively (Ohsawa et al., 2009). Consequently, the native promoter region of the γ-PGA production operon *pgsBCA* was replaced with the xylose-inducible promoter *P_xyl_*. Degradation of the inductor xylose was further inhibited by deletion of *xylAB*, as reported previously (Völker et al., 2025). This yielded the initial production strain *B. subtilis* PG10, referred to as “wild type” in this work. Plasmids required for the subsequent integration or deletion of genes were constructed with the NEB HiFi DNA Assembly Kit (NEB, Frankfurt am Main, Germany) following the manufacturer’s instructions. For the integration, deletion or replacement of genes, homologous target sequences (TS) with a size of roughly 700 bp derived from the up- and downstream sequences of the target locus were amplified and assembled into the linearized pJOE-8739 vector backbone. The following genome engineering of *B. subtilis* by markerless, homologous recombination followed the protocol established by (Wenzel and Altenbuchner, 2015). In brief, plasmids for genome engineering were initially constructed in *E. coli* DH5 α, verified by colony PCR, sequenced and then transformed into *E. coli* JM101 by heat shock transformation. The plasmids obtained from this strain were subsequently transformed into *B. subtilis* using the Paris method (Harwood and Cutting, 1990). Successful transformants were selected on LB agar plates containing spectinomycin. Selected clones were subsequently transferred to LB liquid medium and grown for 8 h at 37°C, where after LB medium containing 0.5 (w/v) mannose was inoculated with 0.02% of the original LB culture. The LB mannose medium was then cultivated for 16 h at 37°C, plated on LB mannose agar plates and emerging colonies were screened for successful plasmid loss by testing for spectinomycin sensitivity. Spectinomycin-sensitive clones were then analyzed by colony PCR to verify gene deletion/integration. For genome engineering of strains *B. subtilis* OF4, 4.2, 5 and 5.2, the protocol was slightly altered to compensate for significantly reduced growth on sole LB medium. In this case, LB medium was complemented with 5 g L^-1^ glucose. Also, the incubation period for the plasmid loss was increased from 8 to 24 and from 16 to 32 h. The gene encoding the Pep carboxylase (*ppc*) was amplified directly from the genome of *C. glutamicum* ATCC 13032 and integrated into the *melA* locus under control of the promoter *P43* or into the *amyE* locus under control of the *Pveg* promoter, reducing genetic redundancy. The hyperactive variant of the glutamate synthase regulator *gltC*_T99A_ was generously provided by Prof. Fabian M. Commichau and integrated into the *odhAB* locus under the control of its native promoter region.

### 2.3 Shake flask cultivation

All presented cultivations were conducted as described in prior work (Völker et al., 2025). Briefly, liquid LB precultures supplemented with 5 g L^-1^ of glucose were prepared in 500 mL unbaffled flasks with a 50 mL working volume. These were inoculated with fresh biomass and incubated for 16 hours at 37 °C with agitation at 200 rpm (25 mm orbital diameter; Infors Multitron, Bottmingen, Switzerland). The resulting precultures were then used to inoculate the main cultures to an initial optical density at 600 nm (OD_600_) of 0.1, determined using an Ultrospec 10 cell density meter (Amersham Bioscience, Little Chalfont, UK). For the main cultures, a modified version of the commonly used Medium E (ME) or mineral salts medium (MSM) was used. Modified medium E contained (per liter): 40 g glycerol, 15 g citrate, 20 g L-glutamate, 10 g NH₄Cl, 0.5 g K₂HPO₄, 0.5 g MgSO₄×7H₂O, 0.04 g FeCl₃×6H₂O, 0.15 g CaCl₂×2H₂O, and 0.104 g MnSO₄×H₂O (Leonard et al., 1958). MSM medium contained (per liter): 40 g glucose, 11.64 g K_2_HPO_4_, 4.89 g NaH_2_PO_4_, 10 g (NH_4_)_2_SO_4_, 0.1 g MgCl_2_×6H_2_O, 0.01 g EDTA, 0.002 g ZnSO_4_×7H_2_O, 0.001 g CaCl_2_×2H_2_O, 0.005 g FeSO_4_×7H_2_O, 0.0002 g Na_2_MoO_4_×2H_2_O, 0.0002 g CuSO_4_×5H_2_O and 0.0004 g CoCl_2_×6H_2_O (Hartmans et al., 1989). Normal MSM medium was supplemented with 0.0001 g L^-1^ MnCl_2_×2H_2_O, whereas adjusted MSM medium contained 0.104 g L^-1^ MnSO₄×H₂O. Induction of the *pgsBCA* operon by *P_xyl_* was ensured by addition of 30 g L^-1^ xylose at the beginning of the experiment. MSM medium was additionally buffered with 200/250/300 mM PIPES buffer. The pH of medium E and MSM medium was adjusted to 7.4 using 10M NaOH. Unless stated otherwise, cultures were then cultivated at 37°C while shaking at 300 rpm (50 mm shaking diameter, Infors Multitron, Bottmingen, Switzerland).

### 2.4 Determination of CO_2_ production and O_2_ consumption

The amount of produced CO_2_ and consumed O_2_ in the off gas of microbial cultures was assessed using the transfer rate online measurement (TOM) shaker (Adolf Kühner AG, Birsfelden, Switzerland). For this, main cultures in adjusted MSM medium were prepared as described in 2.3. The cultures were cultivated at 37°C and shaken at 300 rpm with a shaking diameter of 50 mm. The integral of the respective oxygen transfer rate (OTR) and carbon dioxide transfer rate (CTR) curves was then used to obtain the total amount of consumed O_2_ or produced CO_2_ (Mann et al., 2021).

### 2.5 Calculation of physiological parameters

To assess the max. glucose uptake rate or γ-PGA production rate, the concentration difference between neighboring individual data points was calculated and divided by the time passed. The highest rate determined for the available data points then yielded the max. uptake/production rate obtained in this work. In this window, we further calculated the biomass specific glucose uptake rate according to Takors (2021).

### 2.6 Tracer experiment and ^13^C labeling analysis using GC-MS

Strains were grown for 16 h in a LB + 5 g L^-1^ glucose preculture. Cells were harvested by centrifugation (20 min, 16,000x*g*) during exponential growth, washed once with 0.9% NaCl and used to inoculate the main tracer culture to a starting OD_600_ of 0.05. Natural glucose in the minimal medium was replaced by [1-^13^C] glucose (Kohlstedt et al., 2014). Physiological parameters were collected in a parallel cultivation with biological triplicates and non-labelled glucose (see Tab. S1). During exponential growth (OD_600_ 2, 4, 6), culture broth equivalent to 1 mg of CDW (according to the previously determined OD_600_-to-CDW correlation, see Fig. S2) was collected to extract labeling patterns from protein-bound amino acids and 2.5 mg of CDW were collected to analyze the labeling in cellular carbohydrates (here ribose, glucose and glucosamine). Cells were pelleted by centrifugation (5 min, 4 °C, 16,000x*g*), washed twice with MQ water and stored at -20 °C until further processing. Samples were dried, hydrolyzed, derivatized, and analyzed in selected ion monitoring (SIM) mode via GC-MS as described previously (Kohlstedt and Wittmann, 2019). In total, 36 amino acid and sugar fragments yielded mass isotopomer distributions (MIDs) as input for flux estimation.

### 2.7 Metabolic flux analysis and statistical evaluation

A metabolic network model comprising 100 stoichiometric reactions covering central carbon metabolism (Tab. S2) was reconstructed based on previously published *Bacillus* flux models (He et al., 2019; Kohlstedt et al., 2014; Tännler et al., 2008). Metabolic flux calculation was performed using OpenFLUX (Quek et al., 2009). Experimental mass isotopomer distributions (MIDs) were corrected automatically for the natural abundance of isotopes using the integrated algorithm implemented in OpenFLUX. The metabolic model was further constrained using experimentally determined yields of γ-PGA, acetate, acetoin, lactate, and 2,3-butanediol (Tab. S1), as well as anabolic fluxes toward biomass synthesis (Tab. S3), which were estimated from biomass composition and measured biomass yields. Lower and upper bounds for these constrained fluxes were set using the preSolver function of OpenFLUX (Lehnen et al., 2017), allowing a variation of ±10%. A measurement error of 0.3 mol% was assumed for the GC-MS-derived labelling data (Long et al., 2016). The optimal set of free fluxes was identified by minimising the deviation between experimentally determined and simulated labelling patterns (Tab. S4), and these values were considered to represent the best estimates of the *in vivo* intracellular fluxes (Tab. S5). Flux uncertainty and parameter sensitivity were assessed by Monte Carlo analysis, yielding 95% confidence intervals for all estimated fluxes.

### 2.8 Redox and energy balancing

The estimated intracellular fluxes were used to calculate NAD(P)H and ATP balances. Redox cofactor generation rates (NADPH, NADH, and FADH_2_) were determined from the corresponding cofactor-producing fluxes while accounting for the *in vivo* cofactor specificities of enzymes (Fuhrer and Sauer, 2009; Lerondel et al., 2006). The anabolic NADPH demand was estimated based on biomass composition data (Dauner and Sauer, 2001) and the experimentally determined specific growth rates (Tab. S1). For the cellular energy balance, ATP production via substrate-level phosphorylation (SLP) was calculated by summing ATP-generating fluxes and subtracting ATP-consuming fluxes within the central carbon metabolism network, including PGA biosynthesis. For γ-PGA synthesis, the consumption of 1 mol ATP for ammonium assimilation and 1 mol ATP for polymerisation per mole of glutamate incorporated was assumed. ATP generation via oxidative phosphorylation in the electron transport chain (ETC) was estimated from NADH and FADH_2_ oxidation using a P/O ratio of 1 (Sauer and Bailey, 1999).

The anabolic ATP requirement was calculated from the stoichiometric energy demand for biomass synthesis, using biomass composition data (Dauner and Sauer, 2001) and the measured specific growth rates (Tab. S1). In addition, ATP consumption for non-growth-associated maintenance (NGAM) was included in the overall energy balance (Tännler et al., 2008).

### 2.9 Quantification of intracellular amino acids

The analysis was carried out as described previously (Bolten et al., 2007). Strains were cultivated as described in 2.3 but with 20 g L^-1^ of glucose as carbon source. Further, main cultures were inoculated to a starting OD_600_ of 0.05 and shaken at 230 rpm (25 mm shaking diameter). When the cultures reached an OD_600_ of 1, 3, and 5, culture volume equivalent to 1.2 mg of CDW was collected with a 5 mL plastic syringe and transferred to a filter (cellulose nitrate, 0.2 µm pore size, 25 mm, Sartorius, Göttingen, Germany) for rapid vacuum filtration. The biomass was subsequently washed at room temperature with 15 mL of cultivation medium without carbon source. Afterwards, the filters were transferred into 2 mL of a 200 µM α-aminobutyrate solution, which serves as internal standard, and incubated at 100 °C for 15 min. The resulting extracts were cooled on ice, transferred to 2 mL tubes and centrifuged to precipitate cell debris. Cytosolic amino acids were quantified by HPLC after pre-column derivatisation using *o*-phthalaldehyde (OPA) and 9-fluorenylmethyl chloroformate (FMOC), followed by fluorescence detection, as described previously (Aguilar Suárez et al., 2024). The experiment was performed in triplicates and technical duplicates were collected for each replicate during the sampling process.

### 2.10 γ-PGA quantification with CTAB

Quantitative analysis of γ-PGA titers from bacterial cultures was performed using a cetyltrimethylammonium bromide (CTAB)-based turbidity assay as described in prior work (Halmschlag et al., 2023). Briefly, culture samples were diluted 1:10 and centrifuged for 30 min at 14.000x*g* to remove cells. γ-PGA was then precipitated from cleared culture supernatants by addition of 3x the sample volume of ice cold 99% ethanol. The precipitate was collected by centrifugation, dried and resuspended in distilled water. Afterwards, samples were appropriately diluted to fit into the linear detection range of the assay and mixed in equal portions with a 0.07 M CTAB solution prepared in 2% (w/v) NaOH. The emerging turbidity change was assessed at 400 nm, using a Synergy Mx monochromator-based multimode microplate reader controlled by Gen5 software (both from BioTek Instruments, Inc., Winooski, VT, USA).

### 2.11 Quantification of carbon sources and by-products

The concentration of glucose and the main by-products acetate, acetoin, lactate, and 2,3-butanediol was determined by high-performance liquid chromatography (HPLC-UV/RI) as described previously (Völker et al., 2025). The analysis was carried out using a DIONEX UltiMate 3000 HPLC System (Thermo Scientific, Waltham, MA, USA) equipped with a Metab-AAC column (300 × 7.8 mm, ISERA, Düren, Germany). A 5 mM H₂SO₄ solution was employed as the mobile phase, running isocratically at a flow rate of 0.6 mL/min and maintained at 65°C for 23 minutes. Before injection, the cell-free supernatant was filtered using Rotilabo® syringe filters (0.22 µm, CA), and a 5 µL sample was introduced into the system. Metabolite detection and quantification was achieved with a DIONEX UltiMate 3000 Variable Wavelength Detector set at 210 nm, complemented by a RefractoMax 521 RI detector (Thermo Scientific, Waltham, MA, USA). Metabolite concentrations were determined by comparing the results with calibration curves generated from respective standards.

## 3. Results and discussion

### 3.1 A pyruvate bypass rearranges glycolytic flux to reduce by-product formation

In the first step of our work, we aimed to lower by-product formation by disrupting glycolytic flux towards pyruvate by deleting *pykA*, encoding the sole pyruvate kinase of *B. subtilis* 168. Previous studies have revealed that a subsequent PEP overaccumulation impairs transcription of glycolytic genes, thereby stalling glycolytic flux towards the TCA cycle (Gu et al., 2019). To prevent PEP accumulation, we first expressed the PEP carboxylase gene *ppc* from *C. glutamicum* in our host prior to *pykA* deletion to allow the strain to redirect accumulating PEP towards OAA. This yielded mutant *B. subtilis* **OF1**. Deletion of *pykA* in the OF1 mutant yielded strain *B. subtilis* **OF2**. The generated mutants were cultivated in a common γ-PGA production setup based on Medium E, which mainly consists of glycerol, citrate, and glutamate. Notably, the presence of citrate/glutamate enables pyruvate regeneration through gluconeogenesis via malic enzymes (MleA, MalS, MaeA, YtsJ) or the oxaloacetate decarboxylase (YisK), even in the absence of PykA.

The growth behavior of mutant OF1 was hardly affected by introduction of *ppc* when compared to the wild type (Fig. 2a). However, we observed that the deletion of *pykA* resulted in a remarkable growth defect, likely due to the reduced efficiency of carbon entry into the TCA cycle (Fig. 2a). To test this, a second copy of *ppc* was integrated into the *amyE* locus of *B. subtilis*, yielding strain *B. subtilis* **OF3**. Since the first copy was already equipped with the strong, constitutive P43 promoter, integration of a second copy was favored over a promoter exchange. The additional *ppc* copy restored growth to near wild-type levels (Fig. 2a), whereupon we decided to continue our work with strain OF3. Offline cultivation of the wild type and mutant OF3 showed only minimal differences in OD_600_ and pH development (Fig. 2b). Interestingly, no significant difference in γ-PGA titer (Fig. 2d) and by-product accumulation (Fig. 2e, f) was observed among the newly generated mutant OF3 and the wild type strain in Medium E. To verify the functionality of the introduced bypass, we cultivated strain OF3 in the presence of malate instead of citrate as additional organic acid, as presence of a high malate concentration triggers a strong feedback inhibition on the catalytic activity of Ppc (O’Regan et al., 1989). Addition of malate indeed reduced growth (Fig. 2c) and γ-PGA production (Fig. 2d) of OF3 severely as compared to the wild type, indicating that the activity of Ppc is required to maintain sufficient growth and productivity in a *pykA* deletion mutant (Koendjbiharie et al., 2021). Despite successful rerouting of PEP towards OAA, high levels of overflow metabolites accumulated upon cultivation in Medium E. This likely results from the complex medium composition and the presence of external glutamate. Recent transcriptomic and proteomic analyses of glutamate-dependent γ-PGA production processes showed that glutamate supplementation reduces expression of several TCA cycle genes while promoting pyruvate-derived overflow metabolism (Wu et al., 2025). Together with previous observations that glycerol is mainly converted into biomass and overflow metabolites in citrate/glutamate-containing media (Völker et al., 2025), these findings suggest that external glutamate reduces carbon withdrawal from the TCA cycle for *de novo* glutamate synthesis, thereby masking potential effects of the engineered flux rearrangement. We therefore shifted subsequent experiments to a defined minimal medium.

**Figure 1:**
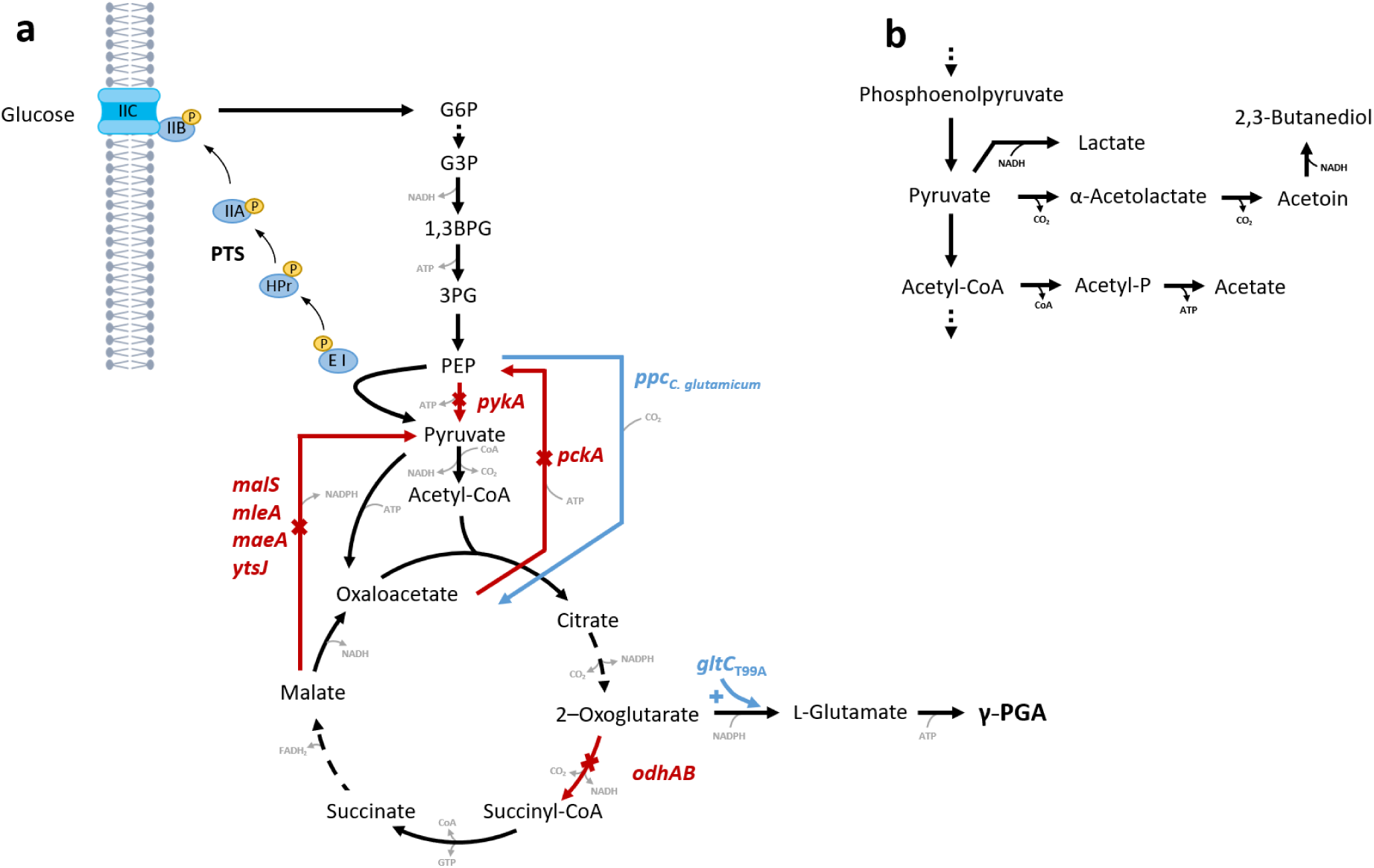
Carbon flux rerouting strategy to minimize pyruvate overflow and by-product accumulation. Under high glucose conditions, *B. subtilis* uses overflow metabolism to convert accumulating pyruvate into by-products, such as acetate, lactate, acetoin and 2,3-butanediol. We here present an engineering strategy, aiming to redirect carbon flow by introduction of an alternative carbon entry route into the TCA cycle (a). Deletions are shown in red, while insertions are shown in blue. The phosphoenolpyruvate carboxylase (*ppc*) from *C. glutamicum* was heterologously expressed in a pyruvate kinase (*pykA*) deletion strain, thereby limiting pyruvate accumulation. With glucose as carbon source, the phosphotransferase system (PTS) with its five subunits (EI, HPr, EIIA, EIIB, EIIC) generates pyruvate from PEP. Genes encoding for enzymes of gluconeogenetic reactions such as the PEP carboxykinase (*pckA*) and the genes of the four malic enzymes (*malS*, *mleA*, *maeA*, *ytsJ*), whos gene products can decarboxylate both malate and oxaloacetate to pyruvate, were successively deleted. Lastly, the operon encoding for the 2-oxoglutarate dehydrogenase complex (*odhAB*) was replaced by a hyperactive variant of the glutamate synthase regulator GltC. Biosynthesis pathways of the main by-products accumulated by *B. subtilis* under excess glucose are shown in (b).

**Figure 2:**
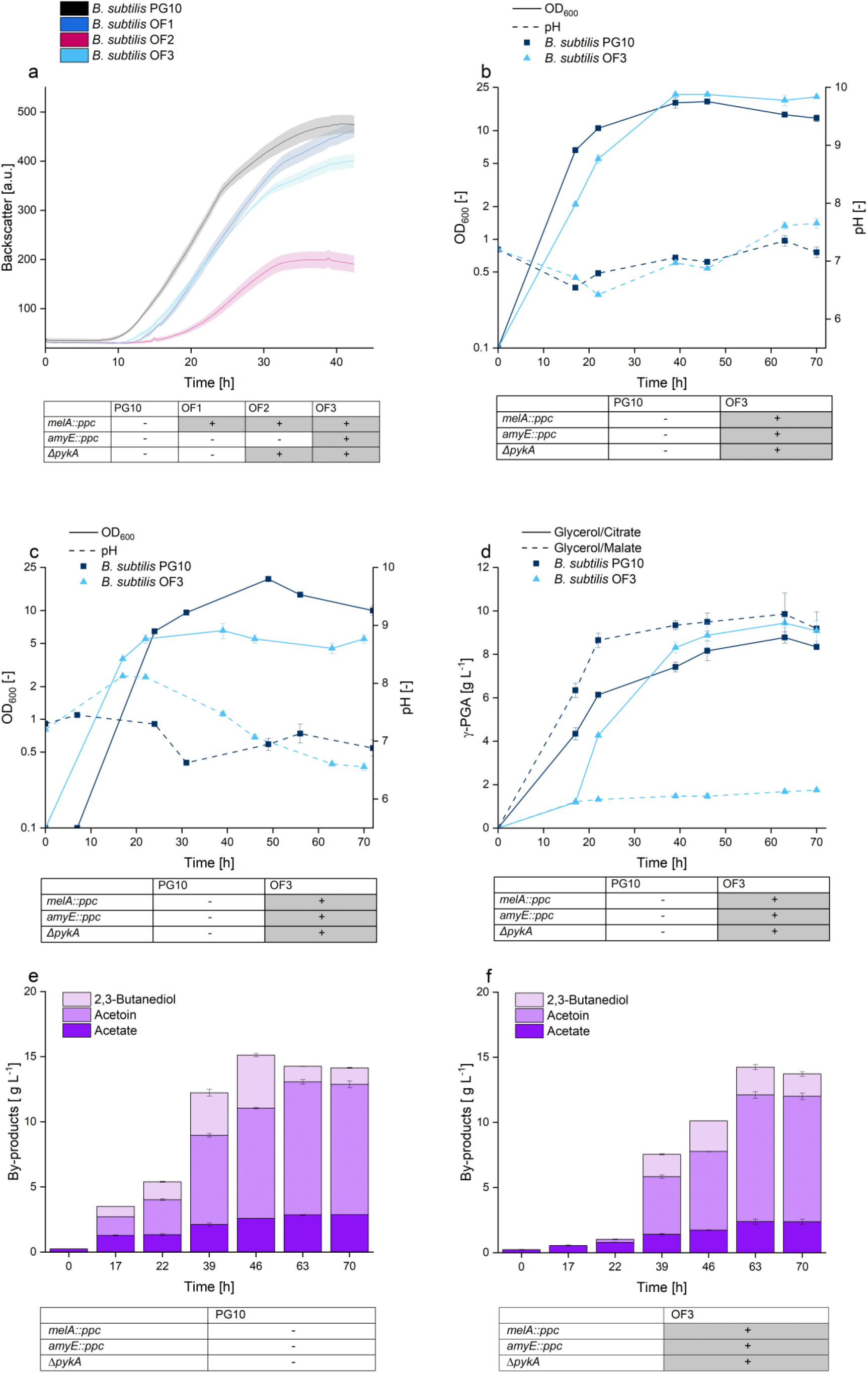
Quantitative physiology of the parental γ-PGA producer *B. subtilis* PG10 and mutants with a heterologous carbon entry route into the TCA cycle. The growth of the parental γ-PGA producer *B. subtilis* PG10 and the mutants OF1, OF2 and OF3 was assessed in ME by online analytics (CGQ, a). The offline growth and pH development of *B. subtilis* PG10 and OF3 in ME with glycerol (40 g L^-1^) and citrate (15 g L^-1^) (b) or glycerol (40 g L^-1^) and malate (15 g L^-1^) as carbon sources (c) is shown. Also, the γ-PGA production in both conditions is depicted (d). By-products accumulated by *B. subtilis* PG10 (e) and strain OF3 (f) are shown. Strain genotypes are indicated below each graph, a (+) indicates that the genotype was changed following the description, (-) implies no genotype changes. The experiment was performed in 500 mL shake flasks with a 50 mL filling volume (a) or 250 mL shake flasks with a 30 mL filling volume (b-f), while shaking at 250 rpm (50 mm shaking diameter).

The mineral salts medium (MSM) is based on glucose as sole carbon source, a phosphate buffer, ammonium sulfate as nitrogen source and trace elements. Previously, glycerol could be used as carbon source, as pyruvate was likely regenerated by gluconeogenesis from either citrate or glutamate (Fig. 2). Notably, none of the so far presented *pykA* deletion mutants grew in MSM with glycerol as sole carbon source due to the lack of an additional substrate for gluconeogenesis (data not shown). Growth was restored when glucose was provided as the sole carbon source instead, likely due to the activity of the phosphotransferase system (PTS), which converts PEP to pyruvate during glucose uptake (Deutscher et al., 2014).

Comparing the composition of Medium E and MSM, a significant difference in the concentration of manganese was detected (1 mg L^-1^ vs. 104 mg L^-1^). Manganese availability has been shown to influence γ-PGA synthesis in *B. subtilis*, as it serves as a co-factor of part of the γ-PGA synthetase PgsBCA (Cromwick and Gross, 1995). We consequently adjusted the manganese concentration in the standard MSM to match the concentration in Medium E. The adjustment slightly affected growth of the wild type, without influencing the pH development or the final OD_600_ reached by the strain (Fig. 3a). Notably, the maximum γ-PGA titer obtained in this adjusted MSM surpassed the titer obtained from standard MSM by 38% (5.2 g L^-1^ vs. 7.2 g L^-1^, Fig. 3 c) with little influence on the glucose uptake rate (10.6 mmol L^-1^ h^-1^ vs 10.8 mmol L^-1^ h^-1^, Fig. 3b) and specific glucose uptake rate (5.6 mmol g_CDW_^-1^ h^-1^ vs. 5.2 mmol g_CDW_^-1^ h^-1^). Increasing the Mn concentration also increased the product yield throughout the whole cultivation, with a maximum product yield of 0.33 g_γ-PGA_ g_glucose_ ^-1^ in adjusted MSM, as compared to 0.24 g_γ-PGA_ g_glucose_ ^-1^ in standard MSM (Fig. 3d).

**Figure 3:**
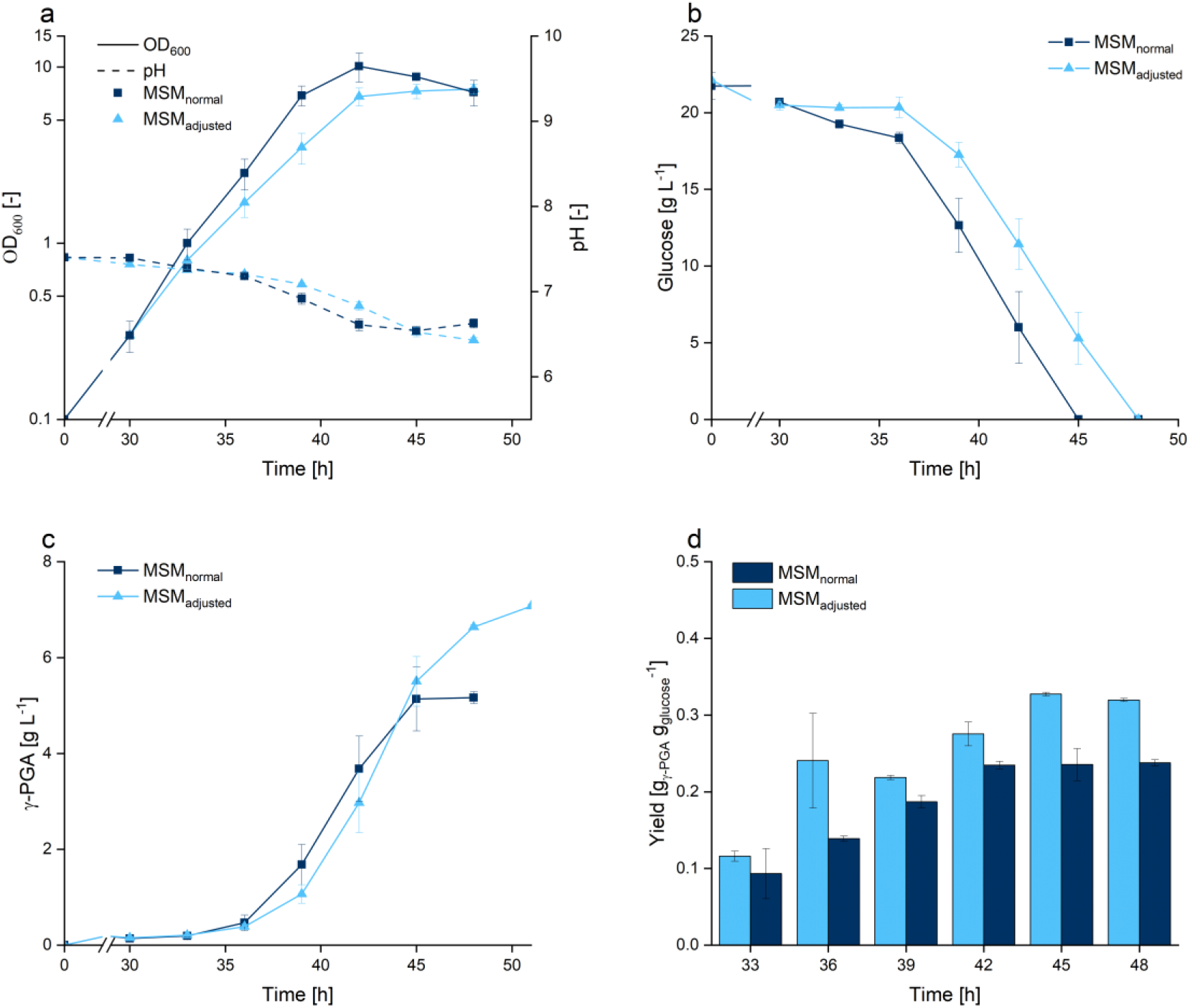
Quantitative physiology of *B. subtilis* PG10 in standard MSM and MSM with increased manganese content. The parental γ-PGA producer *B. subtilis* PG10 was cultivated in standard MSM and in an adjusted version with increased manganese concentration. 200 mM PIPES (pH 7.4) was added as buffer. The growth- and pH development (a), glucose consumption (b), γ-PGA formation (c) and product yield in g_γ-PGA_ g_glucose_^-1^ (d) is shown.

This confirms the adjusted MSM as a suitable setup for γ-PGA synthesis with *B. subtilis* and highlights the importance of manganese for the synthesis of the biopolymer.

### 3.2 Constraining gluconeogenesis reduces the accumulation of overflow metabolites in a Δ*pykA* mutant

The presented strategy bypasses pyruvate accumulation by carbon flux re-routing to OAA, which can be converted to malate by the reverse reaction of the malate dehydrogenase (Mdh) (Koendjbiharie et al., 2021). As presented in literature, high intracellular concentrations of either malate or OAA can trigger activity of gluconeogenetic pathways as response to high TCA cycle flux (Sauer and Eikmanns, 2005). These pathways include the PEP carboxykinase (PckA), which represents a common futile cycle causing ATP loss upon activation, and a total of four malic enzymes (MleA, MalS, MaeA and YtsJ). Although PckA has been reported to reverse its catalytic direction from PEP to OAA in *pykA* deficient *B. subtilis* strains, thereby partially substituting for a missing PEP carboxylase, this reverse reaction proceeds with unfavorable kinetics and is therefore unsuitable to sustain efficient growth (Zamboni et al., 2004). Lerondel et al. (2006) have further reported that the malic enzymes can not only convert malate to pyruvate, but also decarboxylate OAA to pyruvate in addition to the dedicated OAA decarboxylase YisK (Guo et al., 2024). Also, YtsJ is involved in redox balancing by reducing NADP^+^ to NADPH. To ensure that the carbon re-routed by Ppc is maintained in the TCA cycle and not converted back to pyruvate yielding overflow metabolites, we progressively deleted the genes encoding for the enzymes involved in gluconeogenesis. Deletion of *pckA* thereby yielded strain ***B. subtilis* OF4**, and the additional deletions of *mleA*, *malS*, *maeA*, and *ytsJ* yielded strain ***B. subtilis* OF5**.

Growth of the wild type and mutant OF3 was comparable in the adjusted MSM, reaching a final OD_600_ of 11.2 and 11.3 after 45 h of cultivation but with a reduced growth rate of mutant OF3 (Tab. 2, Fig. 4a). In comparison, the mutants OF4 and OF5 grew slower, reaching a final OD of 14.1 and 12.6 after 57 h and 67 h with a further decreased growth rate (Tab. 2). The pH of cultures of the wild type, OF3 and OF4 gradually decreased during the experiment, reaching a final pH of 6.3, 6.1 and 6.2. In the case of mutant OF5, the pH dropped to 5.6 (Fig. 4a). MSM utilizes ammonium sulfate as nitrogen source. Under ammonium excess, which is beneficial for high-level glutamate synthesis, *B. subtilis* consumes the nitrogen source by passive diffusion of uncharged NH_3_. The remaining H^+^ then acidifies the medium, explaining the here observed pH shifts.

**Figure 4:**
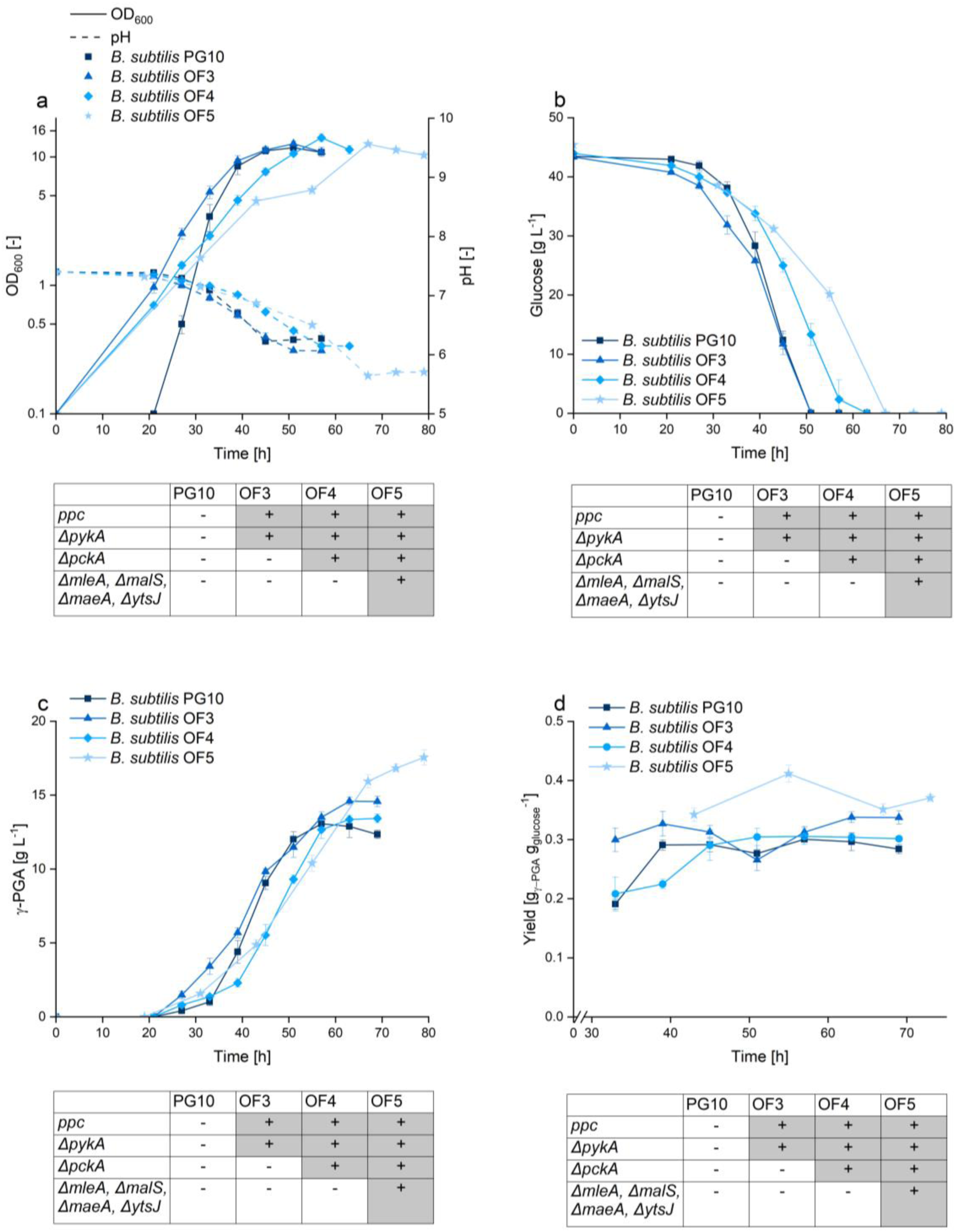
Quantitative physiology of the parental γ-PGA producer *B. subtilis* PG10 and mutants with a heterologous carbon entry route into the TCA cycle and reduced gluconeogenesis. The growth- and pH development (a), glucose consumption (b), γ-PGA formation (c) and product yield (d) of the parental strain *B. subtilis* PG10 and the mutants OF3, OF4 and OF5 is shown. The strains were grown in modified MSM medium with 200 mM PIPES as additional buffer. Strain genotypes are shown below each graph, a (+) indicates that the genotype was changed following the description, (-) implies no genotype changes.

**Table 2:** Key performance indicators of the parental γ-PGA producer *B. subtilis* PG10 and selected mutants. r_S_ and q_S_ were calculated in the window of max. glucose uptake, while r_P_ and q_P_ were calculated in the window of max. product formation. Performance indicators were calculated based on the formulas provided by Takors (2021).

|  | *B. subtilis*<br>PG10 | *B. subtilis*<br>OF3 | *B. subtilis*<br>OF4 | *B. subtilis*<br>OF5 | *B. subtilis*<br>OF4.2 |
| --- | --- | --- | --- | --- | --- |
| $\mu_{\max}$ [h<sup>-1</sup>] | 0.32 ± 0.03 | 0.16 ± 0.01 | 0.12 ± 0.01 | 0.10 ± 0.00 | 0.24 ± 0.00 |
| $Y_{X/S}$ [g<sub>CDW</sub> mol<sub>glucose</sub><sup>-1</sup>] | 19.5 ± 0.6 | 21.0 ± 0.8 | 23.0 ± 1.6 | 19.9 ± 0.8 | 20.3 ± 0.0 |
| Max. glucose uptake<br>window [h] | 39-45 | 39-45 | 51-57 | 55-67 | 18-24 |
| $r_S$ [mmol<sub>glucose</sub> L<sup>-1</sup> h<sup>-1</sup>] | 14.8 ± 1.8 | 13.0 ± 1.6 | 12.3 ± 1.7 | 9.3 ± 0.5 | 20.3 ± 1.7 |
| $q_S$ [mmol<sub>glucose</sub> g<sub>CDW</sub><sup>-1</sup> h<sup>-1</sup>] | 3.8 ± 0.8 | 3.2 ± 0.3 | 2.5 ± 0.5 | 2.7 ± 0.2 | 5.8 ± 0.4 |
| Max. γ-PGA formation<br>window [h] | 39-45 | 39-45 | 45-51 | 43-55 | 18-24 |
| $Y_{P/S}$ [gγ-PGA g<sub>glucose</sub><sup>-1</sup>] | 0.29 ± 0.01 | 0.31 ± 0.01 | 0.30 ± 0.01 | 0.41 ± 0.01 | 0.27 ± 0.02 |
| $r_P$ [gγ-PGA L<sup>-1</sup> h<sup>-1</sup>] | 0.77 ± 0.06 | 0.69 ± 0.04 | 0.63 ± 0.16 | 0.46 ± 0.02 | 1.07 ± 0.11 |
| $q_P$ [gγ-PGA g<sub>CDW</sub><sup>-1</sup> h<sup>-1</sup>] | 0.20 ± 0.03 | 0.17 ± 0.02 | 0.17 ± 0.05 | 0.23 ± 0.02 | 0.31 ± 0.04 |

The differences in growth among the wild type and the mutants were also reflected in the glucose consumption (Tab. 2, Fig. 4b). The max. glucose uptake rate in strains OF4 (12.3 mmol L^-1^ h^-1^) and OF5 (9.3 mmol L^-1^ h^-1^) was concomitantly reduced as compared to the wild type (14.8 mmol L^-1^ h^-1^) and mutant OF3 (13.0 mmol L^-1^ h^-1^). Although increased metabolic interventions led to a delay in strain growth and point of complete glucose consumption (PG10 > OF3 > OF4 > OF5), the biomass yield remained largely consistent across the wild type (19.5 g mol^-1^) and the engineered strains OF3 (21.0 g mol^-1^), OF4 (23.0 g mol^-1^) and OF5 (19.9 g mol^-1^).

Notably, the γ-PGA titer and the accumulation of by-products were significantly affected by the presented rerouting strategy. At the point of complete glucose consumption, the wild type produced a total of 12.0 g L^-1^ γ-PGA (Fig. 4c) and 9.5 g L^-1^ of by products (Fig. 5a, cumulated). The concentrations of both the biopolymer and by-products reached their respective maxima of 13.1 g L⁻¹ for γ-PGA and 9.9 g L⁻¹ for by-products within the next 6 h. Thereafter, a gradual decline in the concentration of both was observed. In *B. subtilis* OF3, glucose was fully depleted after 51 h; however, γ-PGA production continued, reaching a final concentration of 14.6 g L⁻¹ after 63 h (Fig. 4c). In contrast, *B. subtilis* OF4 achieved a maximum γ-PGA titer of 13.4 g L⁻¹. By-product accumulation was effectively reduced in both mutants, with final concentrations of 6.0 g L⁻¹ for OF3 (Fig. 5b) and 5.4 g L⁻¹ for OF4 (Fig. 5c), accounting for a reduction of 39% and 45%.

**Figure 5:**
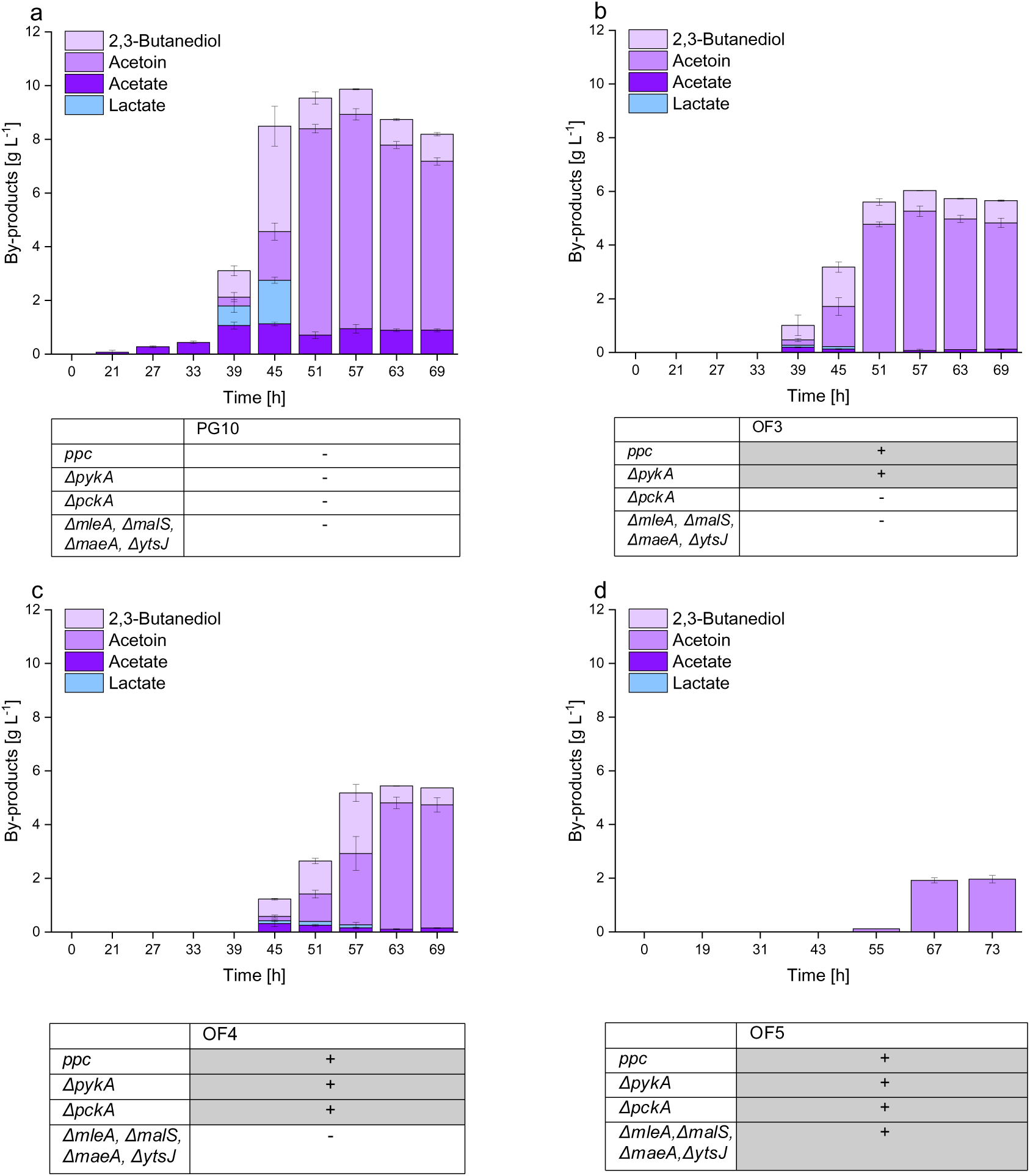
By-products accumulated by the parental γ-PGA producer *B. subtilis* PG10 and mutants with rearranged carbon flux during growth in modified MSM. By-products accumulated by *B. subtilis* PG10 (a), mutant OF3 (b), OF4 (c) and OF5 (d) during the cultivation presented in Figure 4 are shown.

We assumed that the residual by-products were likely produced by gluconeogenesis exerted by malic enzymes. Indeed, accumulation of by-products was nearly eliminated in the mutant *B. subtilis* OF5, lacking the gluconeogenetic pathways encoded by *pckA*, *maeA*, *ytsJ*, *malS* and *mleA*. Only 1.9 g L^-1^ of by-products were detected upon glucose depletion, a decrease of 81% as compared to the wild type (Fig. 5d). Simultaneously, the max. γ-PGA titer rose to 17.6 g L^-1^ in mutant OF5, representing a 35% increase in product titer compared to the wild type strain (Fig. 4c). This is also reflected in the product yield, revealing a max. yield of 0.29 g_γ-PGA_ g_glucose_ ^-1^ for the wild type and 0.41 g_γ-PGA_ g_glucose_ ^-1^ for mutant OF5 (Tab. 2, Fig. 4d).

### 3.3 Deletion of the 2-oxoglutarate dehydrogenase complex increases γ-PGA productivity

While the deletion of gluconeogenetic reactions in mutant OF5 minimized by-product formation and increased γ-PGA yield, it also introduced a significant limitation in process performance, reflected by reduced glucose uptake rates and prolonged cultivation times. We hypothesize that this phenotype is a consequence of reduced metabolic flexibility caused by the removal of the malic enzymes. Preventing the conversion of malate to pyruvate by these deletions may lead to the intracellular accumulation of TCA cycle intermediates, especially malate. As introduced earlier, the catalytic activity of Ppc is feedback-inhibited by elevated intracellular malate concentrations on the protein level, presumably stalling carbon entry into the TCA cycle in mutant OF5 (O’Regan et al., 1989). This would lead to an increase in the intracellular PEP pool. Increased PEP levels have been reported to feedback-regulate the transcriptional levels of the 6-phosphofructokinase *pfkA* and fructose-bisphosphate aldolase *fbaA*, which has been shown to reduce glucose uptake (Gu et al., 2019).

To overcome this limitation, we explored an alternative engineering strategy that retains the malic enzymes to increase the strain’s metabolic flexibility, while controlling carbon flux through the TCA cycle at a different metabolic node. This approach aims to limit carbon flux towards malate formation. By interrupting the TCA cycle downstream of 2-oxoglutarate (2-OX), carbon flux into the lower branch of the TCA cycle is likely reduced, preventing the accumulation of malate. We further hypothesize that this may alleviate Ppc repression, sustain carbon entry into the TCA cycle, and promote a streamlined flux from OAA to 2-OX, while also increasing the pool of 2-OX as a precursor for glutamate synthesis. In this context, Pasotti et al. (2024) have recently reported that successful deletion of genes encoding either the succinyl-CoA synthetase (*sucCD*) or the 2-oxoglutarate dehydrogenase complex (*odhAB*) increases γ-PGA production in *B. subtilis* by enhancing glutamate precursor availability. This observation was also confirmed in earlier studies, highlighting the role of *odhAB* as potential engineering target (Feng et al., 2017; Massaiu et al., 2019). As OdhA catalyzes the irreversible decarboxylation of 2-OX, we decided to interfere on the level of *odhAB* rather than *sucCD* to prevent carbon loss via CO_2_ release. We further hypothesize that while the deletion of *odhAB* may lead to accumulation of 2-OX, the abundance of the glutamate synthase GltAB may be insufficient to effectively withdraw the accumulating precursor. We therefore replaced *odhAB* with a hyperactive version of the transcriptional regulator of the glutamate synthase, GltC. The variant GltC_T99A_ has been shown to drastically increase the expression from the *gltAB* promoter, potentially enabling sufficient transamination of accumulating 2-OX to glutamate (Dormeyer et al., 2019).

In contrast to the previous strategy, this approach favors continuous carbon withdrawal at the 2-OX node to sustain carbon entry into the TCA cycle (*pull*), rather than constraining metabolic flexibility by disrupting gluconeogenetic reactions (*preserve*). The implementation of this strategy (*odhAB*::*gltC*_T99A_) yielded the new mutant ***B. subtilis* OF4.2**. Since a considerable decline in culture pH was observed upon cultivation of mutant OF5 (Fig. 4a), we increased the PIPES buffer concentration in the medium to 250 and 300 mM. With this, we aimed to assess pH-dependent effects on growth and γ-PGA synthesis. While higher buffer concentrations increased the lag phase of strain OF 4.2, likely due to osmotic stress, no significant impact on the overall process parameters were observed compared to buffering with 200 mM PIPES as used in the previous experiments (Fig. 6a). The max. glucose uptake rate of mutant OF4.2 increased considerably (20.3 mmol L^-1^ h^-1^) as compared to the wild type or the previously described mutants (Tab. 2). Analysis of the biomass specific glucose uptake rate for each strain at their respective window of max. glucose uptake rate confirmed that mutant OF4.2 (5.8 mmol g_CDW_^-1^ h^-1^) consumed available glucose faster compared to the wild type (3.8 mmol gCDW^-1^ h^-1^) and the mutants OF3 (3.2 mmol gCDW^-1^ h^-1^), OF4 (2.5 mmol gCDW^-1^ h^-1^) and OF5 (2.7 mmol gCDW^-1^ h^-1^) (Tab. 2). Coherently, faster glucose uptake also resulted in faster γ-PGA formation in strain OF4.2, reaching a max. production rate of 1.07 g_γ-PGA_ L^-1^ h^-1^ (Fig. 6c). This resembles a 39% increase in volumetric productivity as compared to the wild type (0.77 g_γ-PGA_ L^-1^ h^-1^). Interestingly, strain OF4.2 achieved a total γ-PGA titer comparable to that of the wild-type strain upon glucose depletion. However, in contrast to the wild type, the γ-PGA concentration in the cultivation of OF4.2 continued to increase thereafter, ultimately reaching a final titer of 15.0 g L⁻¹. Despite these improvements, mutant OF4.2 accumulated by-products comparable to concentrations observed during the cultivation of the wild type, reaching a cumulative by-product concentration of 9.9 g L^-1^ at the point of glucose depletion. (Fig. 6d). Taken together, the physiological data suggests significant redistributions of intracellular carbon fluxes in the mutants OF4.2 and OF5.

**Figure 6:**
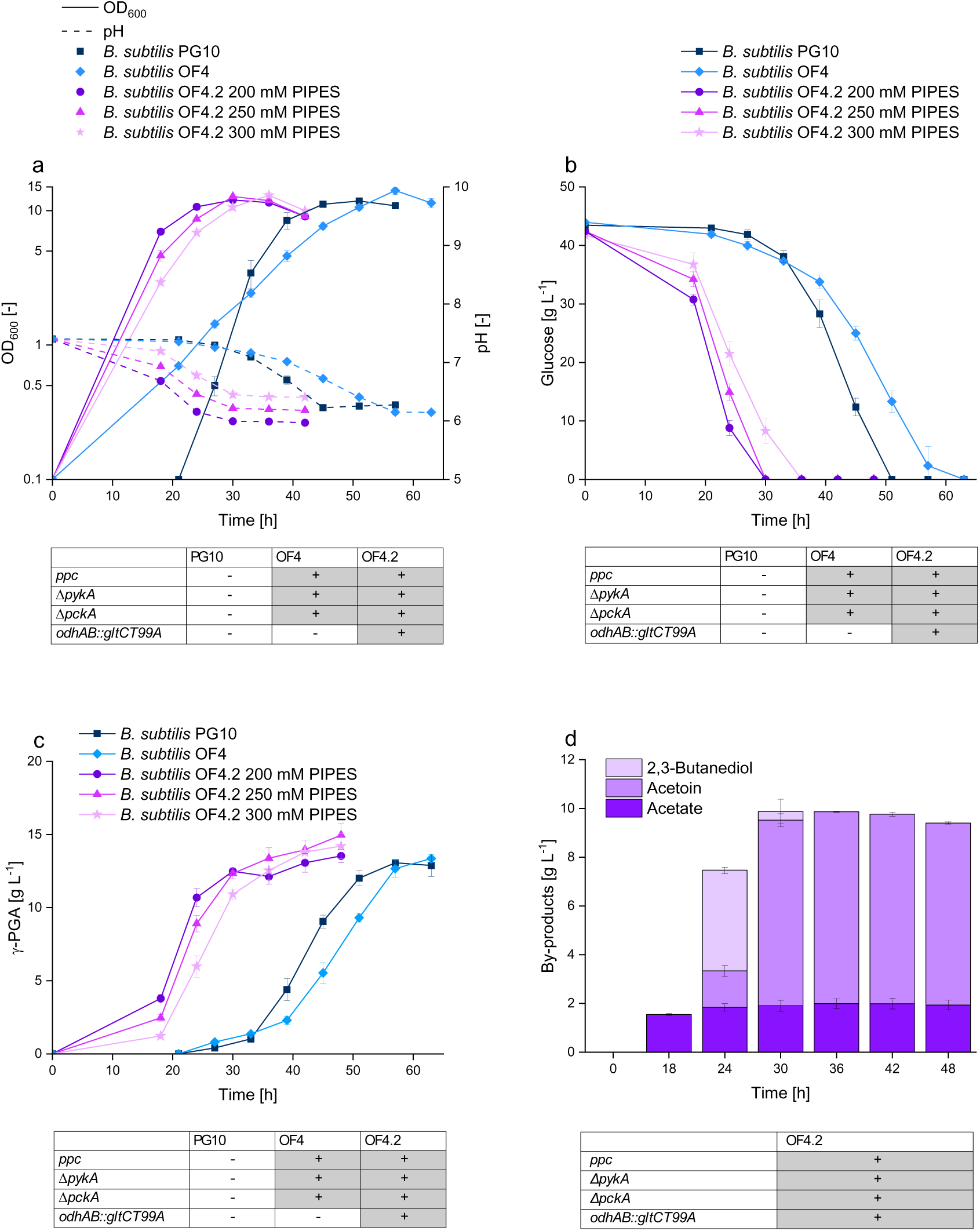
Quantitative physiology of the parental γ-PGA producer *B. subtilis* PG10 and mutants with a heterologous carbon entry route into a disrupted TCA cycle. The growth- and pH development (a), glucose consumption (b) and γ-PGA formation (c) of *B. subtilis* PG10 and the mutants OF4 and OF4.2 is shown. While the strains PG10 and OF4 were cultivated in modified MSM in presence of 200 mM PIPES, strain OF4.2 was cultivated in modified MSM with 200, 250 or 300 mM PIPES. By-products accumulated by mutant OF4.2 in MSM with 200 mM PIPES is shown (d). Strain genotypes are shown below each graph, a (+) indicates that the genotype was changed following the description, (-) implies no genotype changes.

### 3.4 ^13^C metabolic flux analysis and metabolite profiling reveal flux redistribution in mutants OF4.2 and OF5

Since extracellular rates and production kinetics alone are insufficient to draw conclusions on complex flux distributions, ^13^C metabolic flux analysis was performed for the mutants OF4.2, OF5 and the wild type. We report fluxes normalized to the glucose uptake (Fig. 7) and as absolute values (Tab. S5). Unless stated otherwise, the described flux changes were consistent in relative and absolute terms.

**Figure 7:**
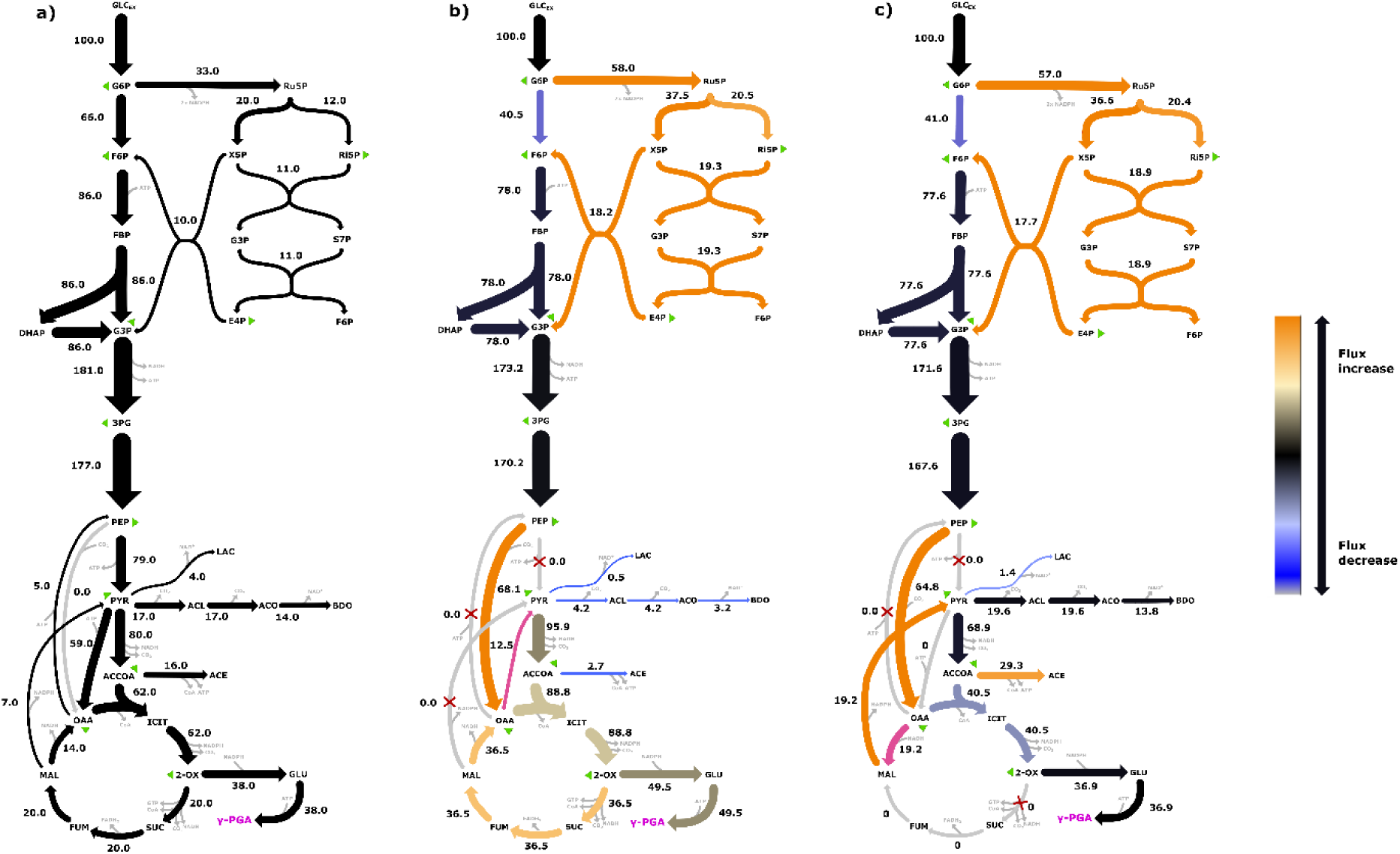
Intracellular carbon fluxes during glucose-based γ-PGA production, revealed by ^13^C metabolic flux analysis. Relative intracellular fluxes of γ-PGA producing *B. subtilis* PG10 (a), mutant OF5 (b) and OF4.2 (c), normalized to the strain-specific glucose uptake rate (set to 100), are shown. Color changes indicate an increased or decreased relative flux as compared to the parental strain. A hypothetical, residual gluconeogenetic flux towards pyruvate in mutant OF5 is indicated in pink, as well as the reverse flux of the malate dehydrogenase. Fluxes towards biomass formation are highlighted by green triangles.

In the wild type, metabolic flux analysis revealed a dominant carbon flux through glycolysis towards pyruvate, with only a moderate fraction of G6P being redirected into the pentose phosphate pathway (PPP). From pyruvate, carbon was mainly channeled into acetyl-CoA and subsequently into the TCA cycle, or channeled towards OAA by activity of the pyruvate carboxylase PycA. The TCA cycle flux downstream of 2-OX was reduced as compared to the upper section of the cycle (OAA to 2-OX), likely due to the steady glutamate demand created by the secretion of γ-PGA (Fig. 7a). Under the tested conditions, flux through gluconeogenetic reactions was low in the wild type strain. In comparison, both mutants OF4.2 and OF5 showed increased flux through the PPP relative to the wild type. The largest flux changes, however, were observed at the branching point between glycolysis and the TCA cycle and in the TCA cycle of the mutants.

In both engineered strains, flux through the disrupted reactions catalyzed by PykA and PckA was set to zero. Carbon entry into the TCA cycle was primarily mediated via Ppc in the mutants, establishing this route as the dominant anaplerotic entry point. In comparison to the wild type, flux through PycA was markedly reduced.

Mutant OF5 exhibited a flux distribution characterized by a strong retention of carbon within the TCA cycle (Fig. 7b). Due to the absence of malic enzyme activity, the conversion of malate to pyruvate was prevented, limiting gluconeogenetic carbon flux towards overflow metabolism. As a result, carbon flux was largely constrained within the TCA cycle, leading to an increased overall TCA cycle flux compared to the wild type. The relative flux towards glutamate synthesis increased, while the absolute flux remained comparable to the wild type. Also, flux from pyruvate into overflow pathways was strongly reduced. A minor flux from OAA towards pyruvate had to be retained in the model for OF5 to achieve a satisfactory fit with the experimental data (Fig. 7b, SI file MIDs). While this flux is required by the model, it does not identify a catalyzing enzyme. Here, the proposed oxaloacetate decarboxylase YisK represents a possible candidate (Guo et al., 2024). The constrained flux distribution is compatible with the proposed accumulation of lower TCA-cycle intermediates in OF5. In particular, increased malate availability could interfere with Ppc-mediated anaplerosis and thereby explain the reduced glucose uptake rate and prolonged process time.

A different carbon flux distribution was observed for mutant OF4.2. The carbon flux downstream of 2-OX was negligible, indicating that carbon loss into the lower branch of the TCA cycle was effectively prevented by *odhAB* deletion (Fig. 7c). Instead, carbon flux originating from OAA was distributed between two major routes. One fraction was directed towards citrate synthesis and subsequently into the upper section of the TCA cycle, while another fraction was channeled towards malate synthesis, likely driven by NADH-dependent, reverse Mdh activity. Malate was subsequently converted to pyruvate via the retained malic enzymes, establishing a cycling of carbon between OAA, malate, and pyruvate. If the retained NADP⁺-dependent malic enzyme YtsJ catalyzes this flux, the cycle couples NADH consumption by reverse Mdh activity to NADPH formation and would constitute a transhydrogenation cycle (Hörl et al., 2021; Lerondel et al., 2006; Rühl et al., 2012). However, the model combines all malic enzyme activities, so the malate-to-pyruvate flux cannot be assigned to a single enzyme and verification of this mechanism will require further testing. From pyruvate, carbon was partially redirected into overflow pathways. Flux towards acetate was increased compared to the wild type, while lactate formation was reduced and flux towards acetoin and 2,3-butanediol remained largely unchanged. Despite these pronounced flux redistributions, the relative flux towards glutamate remained comparable to that of the wild type, while the absolute flux was slightly reduced (Fig. 7c). The metabolic model further predicted a reduced CO₂ formation in mutant OF4.2 compared to the wild type, likely caused by the disruption of the decarboxylation reaction catalyzed by OdhAB. This prediction was compared to measured respiratory activity in shake flasks using the TOM system. Mutant OF4.2 showed both reduced oxygen consumption and lower CO₂ emission compared to the wild type after the available glucose was consumed (see Figure S1). These flux dynamics support the proposed effect of OdhAB disruption, namely the reduction of net carbon flux into the lower TCA cycle while maintaining efficient carbon entry via Ppc. Thus, OF4.2 prevents carbon loss into the lower TCA cycle without enforcing the rigid TCA-cycle confinement observed in OF5. Consistent with previous reports showing that disruption of the TCA cycle increases flux through Ppc (Becker et al., 2009), the presented flux redistribution supports efficient carbon entry into the TCA cycle. At the same time, the presence of the malic enzymes preserves metabolic flexibility by enabling dynamic redistribution of carbon fluxes at the OAA node and may prevent malate accumulation.

This allows the mutant to adjust carbon partitioning between anaplerosis and gluconeogenesis according to cellular demands. Although this flexibility leads to partial carbon dissipation towards by-products, the improved space-time yield of OF4.2 at near-identical wild type titers highlights the potential of the applied engineering strategy. It further raises the question why OF4.2 favors carbon cycling over channeling carbon flux towards glutamate synthesis. We hypothesize that due to strengthened glutamate synthesis in OF4.2, the cell may reach a state of intracellular glutamate saturation due to limited γ-PGA synthesis capacity. This may cause insufficient carbon withdrawal at the 2-OX node, limiting the metabolic “pull” towards glutamate synthesis. To test this hypothesis, we quantified the intracellular concentration of amino acids during different time points of the cultivation. This revealed significant differences between the mutants OF5, OF4.2 and the wild type, consistent with the proposed flux redistribution. In mutant OF4.2, glutamate concentrations remained elevated throughout the cultivation, whereas glutamate levels in both the wild type and OF5 decreased continuously (Fig. 8). This indicates that glutamate supply may exceed glutamate consumption by γ-PGA synthesis in OF4.2. In agreement with this, glutamate-derived amino acids were also affected. Proline accumulated to significantly higher levels in OF4.2 compared to both the wild type and OF5, suggesting that excess glutamate is partially redirected into alternative pathways. At the same time, glutamine levels remained comparatively low in OF4.2, consistent with sustained glutamate synthesis (Fig. 8). Alterations were also observed at the level of amino acids derived from OAA. Aspartate accumulated during later stages of cultivation in both OF4.2 and OF5, while asparagine and threonine showed increased levels particularly in OF4.2. This indicates altered carbon routing at the OAA node, in agreement with anaplerotic carbon entry via Ppc instead of PykA and PycA. In contrast, amino acids derived from other metabolic nodes, including histidine, phenylalanine, tyrosine and isoleucine, remained largely unchanged across all strains and cultivation times, suggesting that the highlighted concentration changes are a specific response to carbon flux remodeling rather than reflecting a global alteration of amino acid biosynthesis (Fig. 8).

**Figure 8:**
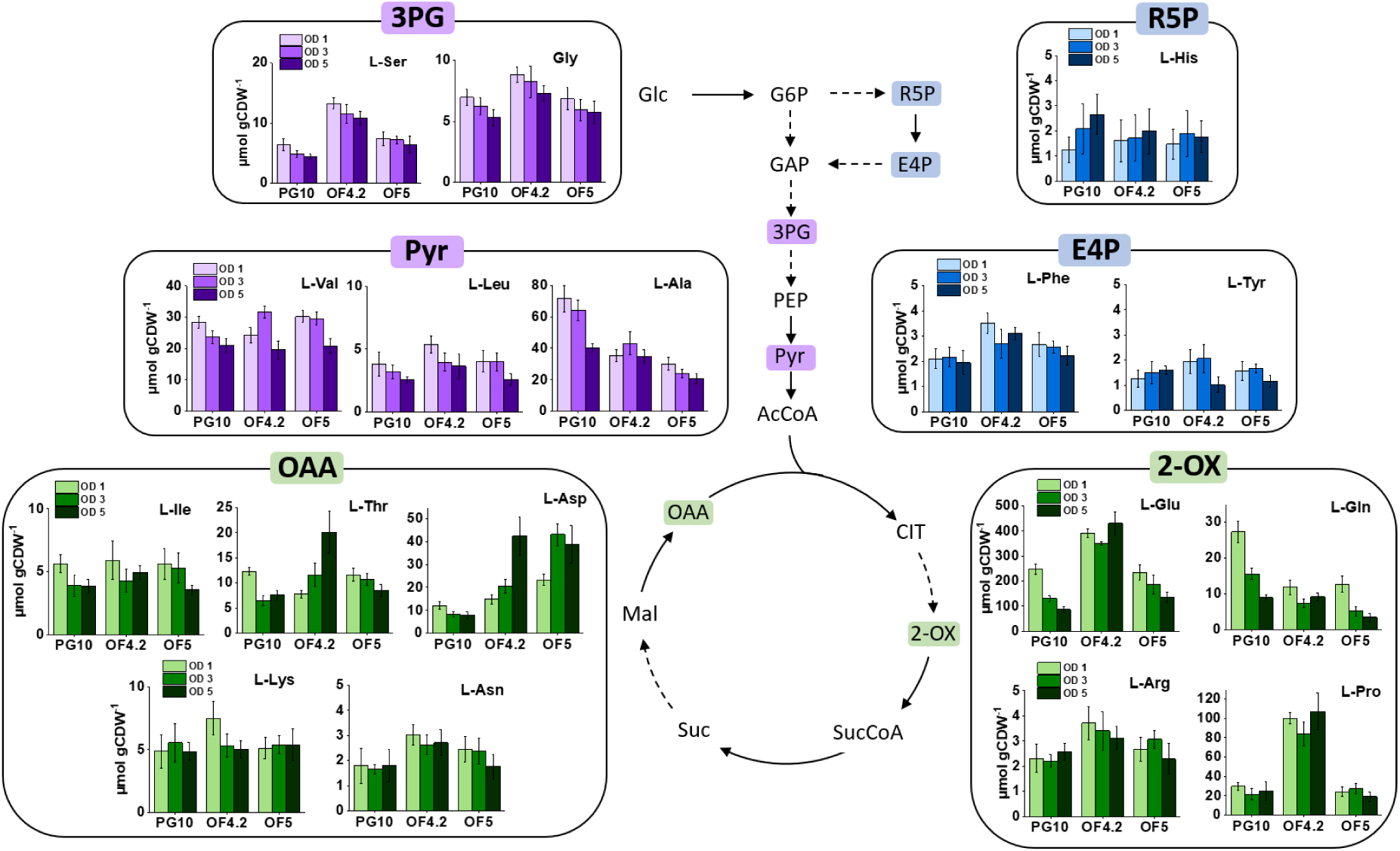
Quantification of intracellular amino acids during γ-PGA synthesis. Samples were collected during growth in adjusted MSM medium when the respective strain had reached an OD of 1, 3 and 5. Cells were subsequently disrupted to enable quantification of intracellular amino acids.

**Figure 9:**
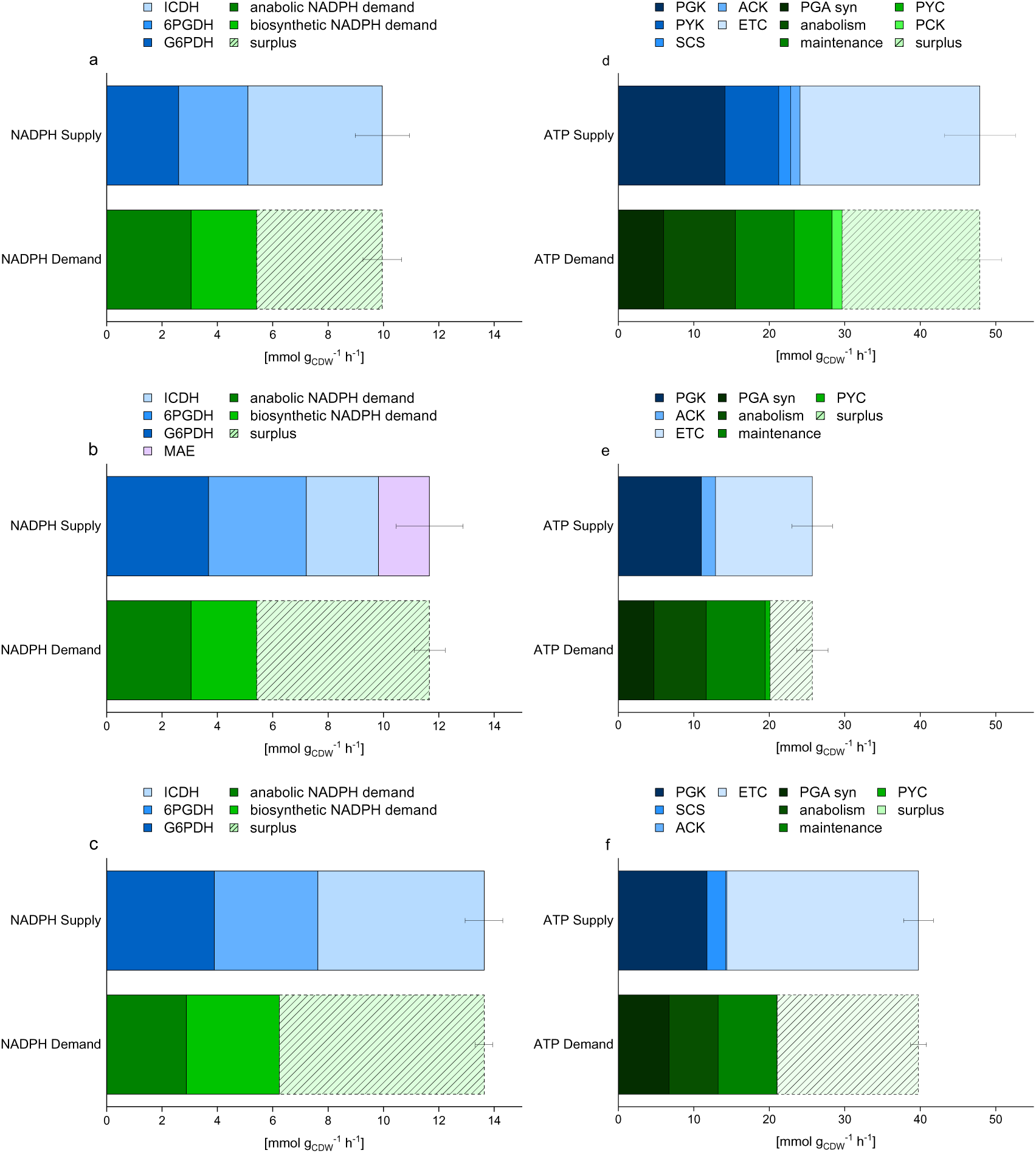
Cellular redox and energy balances of *B. subtilis* during γ-PGA synthesis. The data represent absolute balances for NADPH and ATP in producer *B. subtilis* PG10 (a, d), mutant OF4.2 (b, e) and OF5 (c, f), inferred from the obtained intracellular flux distributions (Figure 7, SI file). Anabolic demand, enzyme cofactor specificities, P/O ratio and maintenance requirements were taken from literature (Dauner and Sauer, 2001; Fuhrer and Sauer, 2009; Sauer and Bailey, 1999; Tännler et al., 2008), considering enzyme cofactor specificities. Abbreviations: ICDH, isocitrate dehydrogenase; 6PGDH, 6-phosphogluconate dehydrogenase; G6PDH, glucose 6-phosphate dehydrogenase; PGK, phosphoglucokinase; SCS, succinyl-CoA synthase; ACK, acetate kinase; ETC, electron transport chain; PGA syn, polyglutamate synthase; PYC, pyruvate carboxylase; PCK, phosphoenolpyruvate carboxykinase; MAE, malic enzyme.

Taken together, the flux analysis and intracellular amino acid profiles highlight that carbon is efficiently routed towards 2-OX and glutamate in mutant OF4.2, while the subsequent conversion into γ-PGA appears to be limiting. This raises the question why the accumulating glutamate is not further polymerized into γ-PGA. One possible explanation is that γ-PGA synthesis capacity becomes limiting, for example due to insufficient *pgsBCA* expression by P_Xyl_ or because the enzyme operates near its maximum capacity. Alternatively, limitations in cellular energy metabolism may restrict γ-PGA synthesis. The carbon cycling between OAA, malate and pyruvate in mutant OF4.2 consumes NADH by Mdh activity, and the interrupted lower TCA cycles forms less ATP and NADH. Both effects may reduce the availability of reducing equivalents and may limit ATP generation. Based on the flux distributions and physiological data, energy and redox balances were calculated. These revealed that ATP supply in mutant OF4.2 was lower compared to both the wild type and mutant OF5. The balance still showed a small ATP surplus in OF4.2, but this surplus was markedly smaller than in the wild type. Given that additional ATP-consuming processes, such as ATP demand for ammonium assimilation, are not fully captured in the model, ATP availability is likely more constrained *in vivo* than predicted. This hypothesis is further supported by the differences in oxygen consumption observed between OF4.2 and the wild type, which agrees with reduced respiratory activity in the mutant strain. As γ-PGA synthesis is an ATP-dependent process, a reduced ATP supply could limit the polymerization of glutamate into γ-PGA. At the same time, increased NADH turnover may lead to rapid regeneration of NAD⁺. An increased availability of NAD⁺ may contribute to increased glycolytic flux and thereby explain the elevated glucose uptake rates in mutant OF4.2. However, the extent of how glycolytic activity is limited by NAD⁺ availability remains speculative.

Moreover, an increased flux through the PPP was observed in both mutants. The considerable intracellular glutamate accumulation observed in OF4.2 (Fig. 8) indicates an increased NADPH demand, for example via the NADPH-dependent glutamate synthase activity, that may exceed the supply from the PPP alone. The elevated PPP flux observed in OF4.2 is consistent with this increased demand, and the proposed transhydrogenation cycle may provide an additional NADPH synthesis route. In mutant OF5, elevated PPP flux instead appears to originate from stalled anaplerotic carbon entry through Ppc rather than from an increased NADPH demand, and may therefore not be linked to a comparable NADPH-generating role.

These observations highlight γ-PGA production in strain OF4.2 as a trade-off between high glucose uptake and production rates and limitations caused by adverse carbon flux and imbalanced redox metabolism. Alleviating these hurdles will require further strain engineering to maintain high carbon throughput and improved ATP availability while reducing carbon cycling. In this context, direct removal of the malic enzymes in OF4.2 was not feasible, as the resulting strains showed detrimental growth defects (data not shown). This suggests that a certain degree of carbon cycling between OAA, malate and pyruvate may be required to maintain cellular fitness under conditions of high carbon flux as presented in mutant OF4.2.

## 4. Conclusion

This study demonstrates that optimizing γ-PGA production in *B. subtilis* is not solely dependent on increasing carbon flux towards glutamate synthesis but requires additional control of redox metabolism and energy generation. This interpretation is supported by earlier studies in *B. licheniformis* where modulation of pentose phosphate pathway activity and cellular energy metabolism was required to improve γ-PGA production, indicating that precursor supply alone is insufficient in scenarios of cofactor or ATP limitation (Cai et al., 2017).

Moreover, the engineering strategies presented in this work show distinct metabolic responses of *B. subtilis* depending on how carbon flux is directed. While constraining carbon flux within the TCA cycle increased product yield by limiting carbon loss due to overflow metabolism in mutant OF5, substrate uptake and product formation rates were limited. In contrast, maintaining high glucose uptake and production rates in mutant OF4.2 supported rapid substrate uptake and product formation. Carbon partitioning at the OAA node shifted towards cycling between OAA, malate and pyruvate. This coincided with limitations at the level of redox cofactor supply and energy metabolism. A similar requirement for a coordinated optimization of carbon flux and redox metabolism has been reported by Gu et al. (2019) while improving GlcNAc production in *B. subtilis*. Further strain engineering should focus on reducing NADH drainage at the OAA/malate/pyruvate node to alleviate possible ATP limitations during γ-PGA synthesis in mutant OF4.2. This may be achieved by the targeted modulation of Mdh activity or the introduction of enzyme variants with altered cofactor specificity.

Together, the presented integration of physiological strain characterization with flux analysis and intracellular metabolite profiling showcases key metabolic constraints at the intersection of glycolysis and the TCA cycle.

## 6. Acknowledgements

We thank Prof. Fabian M. Commichau for the generous provision of the plasmid carrying the hyperactive *gltC*_T99A_ variant.

## 7. Author contributions

FV conceptualized the study, performed the experiments, evaluated the data, prepared the original draft and revised the manuscript. MK performed experiments, evaluated the data and revised the manuscript. BH performed experiments and revised the manuscript. JEM and JSW performed experiments. CW revised the manuscript. LMB conceptualized the study, provided supervision, acquired funding, and revised the manuscript.

## 8. Declarations

The authors have no conflicts of interest to declare. During the preparation of this work the author(s) used OpenAI’s Assistant and Grammarly to edit the language of this manuscript. After using this tool/service, the author(s) reviewed and edited the content as needed and take(s) full responsibility for the content of the published article.

## 9. Data availability

The physiological data and the data of the metabolic flux analysis supporting the findings of this study are available from the corresponding author upon reasonable request.

## References

Aguilar Suárez, R., Kohlstedt, M., Öktem, A., Neef, J., Wu, Y., Ikeda, K., Yoshida, K.I., Altenbuchner, J., Wittmann, C., van Dijl, J.M., 2024. Metabolic profile of the genome-reduced *Bacillus subtilis* strain IIG-Bs-27-39: An attractive chassis for recombinant protein production. ACS Synth. Biol. 13, 2199–2214,

Balogun-Agbaje, O.A., Odeniyi, O.A., Odeniyi, M.A., 2021. Drug delivery applications of poly-γ-glutamic acid. Futur. J. Pharm. Sci. 2021 71 7, 1–10,

Basan, M., Hui, S., Okano, H., Zhang, Z., Shen, Y., Williamson, J.R., Hwa, T., 2015. Overflow metabolism in *E. coli* results from efficient proteome allocation. Nature 528, 99,

Bolten, C.J., Kiefer, P., Letisse, F., Portais, J.C., Wittmann, C., 2007. Sampling for metabolome analysis of microorganisms. Anal. Chem. 79, 3843–3849,

Cai, D., He, P., Lu, X., Zhu, C., Zhu, J., Zhan, Y., Wang, Q., Wen, Z., Chen, S., 2017. A novel approach to improve poly-γ-glutamic acid production by NADPH regeneration in *Bacillus licheniformis* WX-02. Sci. Rep. 7, 43404,

Chen, T., Brul, S., Hugenholtz, J., 2023. Exploring the potential of *Bacillus subtilis* as cell factory for food ingredients and special chemicals. Microb. Cell Fact. 22, 1–12,

Cromwick, A.M., Gross, R.A., 1995. Effects of manganese (II) on *Bacillus licheniformis* ATCC 9945A physiology and γ-poly(glutamic acid) formation. Int. J. Biol. Macromol. 17, 259– 267,

Dauner, M., Sauer, U., 2001. Stoichiometric growth model for riboflavin-producing *Bacillus subtilis*. Biotechnol. Bioeng. 76, 132–143,

Deutscher, J., Moussan, F., Aké, D., Derkaoui, M., Constant Zébré, A., Cao, T.N., Bouraoui, H., Kentache, T., Mokhtari, A., Milohanic, E., Joyet, P., Aké, F.M.D., Derkaoui, M., Zébré, A.C., Cao, T.N., Bouraoui, H., Kentache, T., Mokhtari, A., Milohanic, E., Joyet, P., 2014. The bacterial phosphoenolpyruvate:carbohydrate phosphotransferase system: Regulation by protein phosphorylation and phosphorylation-dependent protein-protein interactions. Microbiol. Mol. Biol. Rev. 78, 231,

Dormeyer, M., Lentes, S., Richts, B., Heermann, R., Ischebeck, T., Commichau, F.M., 2019. Variants of the *Bacillus subtilis* LysR-Type regulator GltC with altered activator and repressor function. Front. Microbiol. 10, 488960,

Ewing, T.A., Nouse, N., van Lint, M., van Haveren, J., Hugenholtz, J., van Es, D.S., 2022. Fermentation for the production of biobased chemicals in a circular economy: a perspective for the period 2022–2050. Green Chem. 24, 6373–6405,

Fan, Z., Cheng, P., Liu, M., Li, D., Liu, G., Zhao, Y., Ding, Z., Chen, F., Wang, B., Tan, X., Wang, Z., Han, J., 2017. Poly(glutamic acid) hydrogels crosslinked via native chemical ligation. New J. Chem. 41, 8656–8662,

Feng, J., Quan, Y., Gu, Y., Liu, F., Huang, X., Shen, H., Dang, Y., Cao, M., Gao, W., Lu, X., Wang, Y., Song, C., Wang, S., 2017. Enhancing poly-γ-glutamic acid production in *Bacillus amyloliquefaciens* by introducing the glutamate synthesis features from *Corynebacterium glutamicum*. Microb. Cell Fact. 16, 1–12,

Fuhrer, T., Sauer, U., 2009. Different biochemical mechanisms ensure network-wide balancing of reducing equivalents in microbial metabolism. J. Bacteriol. 191, 2112–2121,

Gu, Y., Lv, X., Liu, Y., Li, J., Du, G., Chen, J., Rodrigo, L.A., Liu, L., 2019. Synthetic redesign of central carbon and redox metabolism for high yield production of N-acetylglucosamine in *Bacillus subtilis*. Metab. Eng. 51, 59–69,

Guo, T., Sperber, A.M., Krieger, I. V., Duan, Y., Chemelewski, V.R., Sacchettini, J.C., Herman, J.K., 2024. *Bacillus subtilis* YisK possesses oxaloacetate decarboxylase activity and exhibits Mbl-dependent localization. J. Bacteriol. 206,

Halmschlag, B., Hoffmann, K., Hanke, R., Putri, S.P., Fukusaki, E., Büchs, J., Blank, L.M., 2020. Comparison of Isomerase and Weimberg pathway for γ-PGA production from xylose by engineered *Bacillus subtilis*. Front. Bioeng. Biotechnol. 7, 476,

Halmschlag, B., Steurer, X., Putri, S.P., Fukusaki, E., Blank, L.M., 2019. Tailor-made poly-γ-glutamic acid production. Metab. Eng. 55, 239–248,

Halmschlag, B., Völker, F., Hanke, R., Putri, S.P., Fukusaki, E., Büchs, J., Blank, L.M., 2023. Metabolic engineering of *B. subtilis* 168 for increased precursor supply and poly-γ-glutamic acid production. Front. Food Sci. Technol. 3,

Hartmans, S., Smits, J.P., Van der Werf, M.J., Volkering, F., De Bont, J.A.M., 1989. Metabolism of styrene oxide and 2-phenylethanol in the styrene-degrading *Xanthobacter* strain 124X. Appl. Environ. Microbiol. 55, 2850–2855,

Harwood, C.R., Cutting, S.M., 1990. Molecular biological methods for *Bacillus*. Mol. Biol. Methods Bacillus 493–543,

He, P., Wan, N., Cai, D., Hu, S., Chen, Y., Li, S., Chen, S., 2019. ^13^C-Metabolic Flux Analysis reveals the metabolic flux redistribution for enhanced production of poly-γ-glutamic acid in *dlt* over-expressed *Bacillus licheniformis*. Front. Microbiol. 10,

Hörl, M., Fuhrer, T., Zamboni, N., 2021. Bifunctional malic/malolactic enzyme provides a novel mechanism for NADPH-balancing in *Bacillus subtilis*. MBio 12,

Ioannidou, S.M., Pateraki, C., Ladakis, D., Papapostolou, H., Tsakona, M., Vlysidis, A., Kookos, I.K., Koutinas, A., 2020. Sustainable production of bio-based chemicals and polymers via integrated biomass refining and bioprocessing in a circular bioeconomy context. Bioresour. Technol. 307, 123093,

Jiang, Y., Tang, B., Xu, Zongqi, Liu, K., Xu, Zheng, Feng, X., Xu, H., 2016. Improvement of poly-γ-glutamic acid biosynthesis in a moving bed biofilm reactor by *Bacillus subtilis* NX-2. Bioresour. Technol. 218, 360–366,

Kabisch, J., Pratzka, I., Meyer, H., Albrecht, D., Lalk, M., Ehrenreich, A., Schweder, T., 2013. Metabolic engineering of *Bacillus subtilis* for growth on overflow metabolites. Microb. Cell Fact. 12, 72,

Koendjbiharie, J.G., Van Kranenburg, R., Kengen, S.W.M., 2021. The PEP-pyruvate-oxaloacetate node: variation at the heart of metabolism. FEMS Microbiol. Rev. 45, 1–19,

Kohlstedt, M., Sappa, P.K., Meyer, H., Maaß, S., Zaprasis, A., Hoffmann, T., Becker, J., Steil, L., Hecker, M., van Dijl, J.M., Lalk, M., Mäder, U., Stülke, J., Bremer, E., Völker, U., Wittmann, C., 2014. Adaptation of *Bacillus subtilis* carbon core metabolism to simultaneous nutrient limitation and osmotic challenge: a multi-omics perspective. Environ. Microbiol. 16, 1898–1917,

Kohlstedt, M., Wittmann, C., 2019. GC-MS-based ^13^C metabolic flux analysis resolves the parallel and cyclic glucose metabolism of *Pseudomonas putida* KT2440 and *Pseudomonas aeruginosa* PAO1. Metab. Eng. 54, 35–53,

Lehnen, M., Ebert, B.E., Blank, L.M., 2017. A comprehensive evaluation of constraining amino acid biosynthesis in compartmented models for metabolic flux analysis. Metab. Eng. Commun. 5, 34–44,

Leonard, C.G., Housewright, R.D., Thorne, C.B., 1958. Effects of some metallic ions on glutamyl polypeptide synthesis by *Bacillus subtilis*. J. Bacteriol. 76, 499–503,

Lerondel, G., Doan, T., Zamboni, N., Sauer, U., Aymerich, S., 2006. YtsJ has the major physiological role of the four paralogous malic enzyme isoforms in *Bacillus subtilis*. J. Bacteriol. 188, 4727,

Li, Z., Nees, M., Bettenbrock, K., Rinas, U., 2022. Is energy excess the initial trigger of carbon overflow metabolism? Transcriptional network response of carbon-limited *Escherichia coli* to transient carbon excess. Microb. Cell Fact. 21, 1–19,

Long, C.P., Au, J., Gonzalez, J.E., Antoniewicz, M.R., 2016. ^13^C metabolic flux analysis of microbial and mammalian systems is enhanced with GC-MS measurements of glycogen and RNA labeling. Metab. Eng. 38, 65–72,

Ma, W., Liu, Y., Shin, H. dong, Li, J., Chen, J., Du, G., Liu, L., 2018. Metabolic engineering of carbon overflow metabolism of *Bacillus subtilis* for improved N-acetyl-glucosamine production. Bioresour. Technol. 250, 642–649,

Mann, M., Wittke, D., Büchs, J., 2021. Online monitoring applying the anaerobic respiratory monitoring system reveals iron(II) limitation in YTF medium for *Clostridium ljungdahlii*. Eng. Life Sci. 21, 19–28,

Massaiu, I., Pasotti, L., Sonnenschein, N., Rama, E., Cavaletti, M., Magni, P., Calvio, C., Herrgård, M.J., 2019. Integration of enzymatic data in *Bacillus subtilis* genome-scale metabolic model improves phenotype predictions and enables in silico design of poly-γ-glutamic acid production strains. Microb. Cell Fact. 18,

Mesa, J.A., Sierra-Fontalvo, L., Ortegon, K., Gonzalez-Quiroga, A., 2024. Advancing circular bioeconomy: A critical review and assessment of indicators. Sustain. Prod. Consum. 46, 324–342,

Meselson, M., Yuan, R., 1968. DNA restriction enzyme from *E. coli*. Nat. 1968 2175134 217, 1110–1114,

Messing, J., Crea, R., Seeburg, P.H., 1981. A system for shotgun DNA sequencing. Nucleic Acids Res. 9, 309,

Molenaar, D., Van Berlo, R., De Ridder, D., Teusink, B., 2009. Shifts in growth strategies reflect tradeoffs in cellular economics. Mol. Syst. Biol. 5, 323,

Nair, P., Navale, G.R., Dharne, M.S., 2023. Poly-gamma-glutamic acid biopolymer: a sleeping giant with diverse applications and unique opportunities for commercialization. Biomass Convers. Biorefinery 13, 4555,

O’Regan, M., Thierbach, G., Bachmann, B., Villeval, D., Lepage, P., Viret, J.F., Lemoine, Y., 1989. Cloning and nucleotide sequence of the phosphoenolpyruvate carboxylase-coding gene of *Corynebacterium glutamicum* ATCC13032. Gene 77, 237–251,

Ohsawa, T., Tsukahara, K., Ogura, M., 2009. *Bacillus subtilis* response regulator DegU is a direct activator of *pgsB* transcription involved in gamma-poly-glutamic acid synthesis. Biosci. Biotechnol. Biochem. 73, 2096–2102,

Park, S. Bin, Sung, M.H., Uyama, H., Han, D.K., 2021. Poly(glutamic acid): Production, composites, and medical applications of the next-generation biopolymer. Prog. Polym. Sci. 113, 101341,

Pasotti, L., Massaiu, I., Magni, P., Calvio, C., 2024. Metabolic engineering of *Bacillus subtilis* for the production of poly-γ-glutamic acid from glycerol feedstock. Ferment. 10, 319,

Peng, Y., Jiang, B., Zhang, T., Mu, W., Miao, M., Hua, Y., 2015. High-level production of poly(γ-glutamic acid) by a newly isolated glutamate-independent strain, *Bacillus methylotrophicus*. Process Biochem. 50, 329–335,

Quek, L.E., Wittmann, C., Nielsen, L.K., Krömer, J.O., 2009. OpenFLUX: efficient modelling software for ^13^C-based metabolic flux analysis. Microb. Cell Fact. 8,

Rühl, M., Le Coq, D., Aymerich, S., Sauer, U., 2012. ^13^C-flux analysis reveals NADPH-balancing transhydrogenation cycles in stationary phase of nitrogen-starving *Bacillus subtilis*. J. Biol. Chem. 287, 27959,

Sauer, U., Bailey, J.E., 1999. Estimation of P-to-O Ratio in *Bacillus subtilis* and its influence on maximum riboflavin yield. Biotechnol. Bioeng,

Sauer, U., Eikmanns, B.J., 2005. The PEP—pyruvate—oxaloacetate node as the switch point for carbon flux distribution in bacteria. FEMS Microbiol. Rev. 29, 765–794,

Sheldon, R.A., Brady, D., 2022. Green Chemistry, biocatalysis, and the chemical industry of the future. ChemSusChem 15,

Shimizu, K., Matsuoka, Y., 2019. Regulation of glycolytic flux and overflow metabolism depending on the source of energy generation for energy demand. Biotechnol. Adv. 37, 284–305,

Skalski, T., Zając, E., Jędrszczyk, E., Papaj, K., Kohyt, J., Góra, A., Kasprzycka, A., Shytum, D., Skowera, B., Ziernicka-Wojtaszek, A., 2024. Effects of γ-polyglutamic acid on grassland sandy soil properties and plant functional traits exposed to drought stress. Sci. Rep. 14, 3769,

Stülke, J., Grüppen, A., Bramkamp, M., Pelzer, S., 2023. *Bacillus subtilis*, a swiss army knife in science and biotechnology. J. Bacteriol. 205,

Takors, R., 2021. Kommentierte Formelsammlung Bioverfahrenstechnik. Spring. Nat. ISBN 978-3-662-63971-9,

Tännler, S., Decasper, S., Sauer, U., 2008. Maintenance metabolism and carbon fluxes in *Bacillus* species. Microb. Cell Fact. 7,

Völker, F., Hoffmann, K., Halmschlag, B., Maaß, S., Büchs, J., Blank, L.M., 2025. Citrate supplementation modulates medium viscosity and poly-γ-glutamic acid synthesis by engineered *B. subtilis* 168. Eng. Life Sci. 25, e70009,

Wenzel, M., Altenbuchner, J., 2015. Development of a markerless gene deletion system for *Bacillus subtilis* based on the mannose phosphoenolpyruvate-dependent phosphotransferase system. Microbiol. (United Kingdom) 161, 1942–1949,

Wu, C., Gou, Y., Jing, S., Li, W., Ge, F., Li, J., Ren, Y., 2025. Analysis of glutamate-dependent mechanism and optimization of fermentation conditions for poly-gamma-glutamic acid production by *Bacillus subtilis* SCP017-03. PLoS One 20, e0310556,

Yokota, A., Sawada, K., Wada, M., 2017. Boosting anaplerotic reactions by pyruvate kinase gene deletion and phosphoenolpyruvate carboxylase desensitization for glutamic acid and lysine production in *Corynebacterium glutamicum*. Adv. Biochem. Eng. Biotechnol. 159, 181–198,

Yuan, P., Xu, M., Mao, C., Zheng, H., Sun, D., 2023. Dynamically regulating glucose uptake to reduce overflow metabolism with a quorum-sensing circuit for the efficient synthesis of d-pantothenic acid in *Bacillus subtilis*. ACS Synth. Biol. 12, 2983–2995,

Zamboni, N., Maaheimo, H., Szyperski, T., Hohmann, H.P., Sauer, U., 2004. The phosphoenolpyruvate carboxykinase also catalyzes C3 carboxylation at the interface of glycolysis and the TCA cycle of *Bacillus subtilis*. Metab. Eng. 6, 277–284.

