## Supplementary material for "Rewiring central carbon metabolism reveals a trade-off between yield and production rate in poly-γ-glutamic acid synthesis by *Bacillus subtilis*"

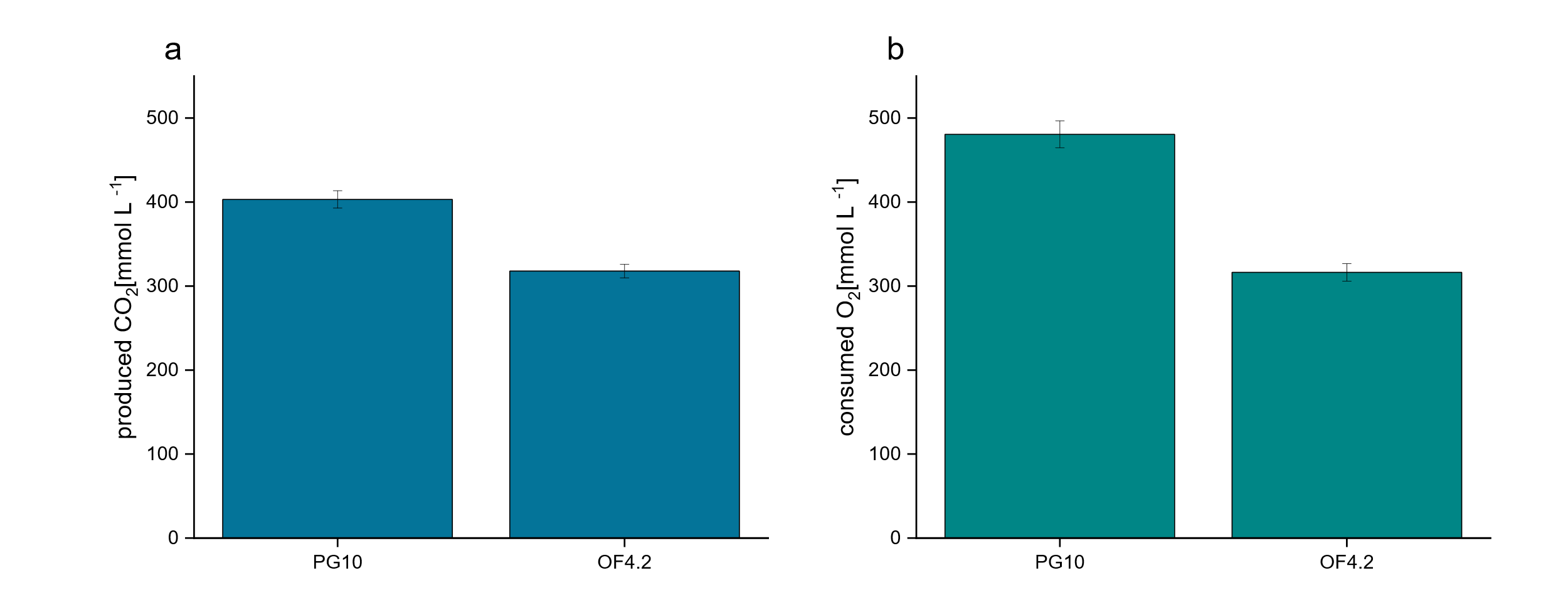

**Figure S1: End-point CO_2_ production and O_2_ consumption of B. subtilis PG10 and mutant OF4.2**

The parental γ-PGA producer B. subtilis PG10 and the mutant strain OF4.2 were cultivated in modified MSM following the same conditions as shown in Figure 4. The total CO_2_ production (a) of the strains and the total amount of consumed O_2_ (b) at the point of complete glucose consumption is shown.

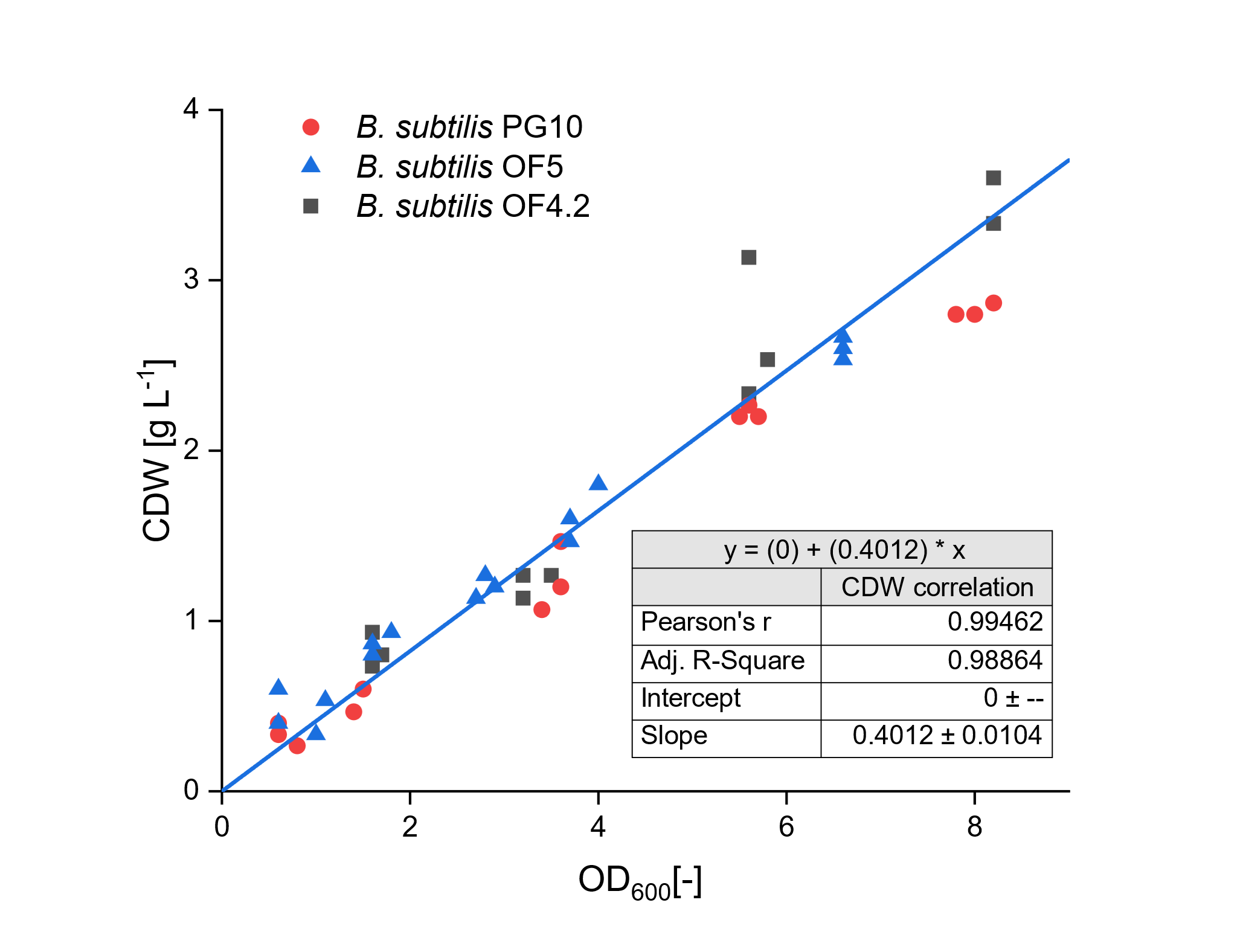

**Figure S2: OD_600_ to CDW correlation of B. subtilis strains**

Strains were grown in modified MSM with 20 g L^-1^ of glucose.

**Table S1: Key physiological parameters of γ-PGA producing B. subtilis strains to constrain the metabolic modeling procedure.**

| **Physiological parameter** | ***B. subtilis* PG10** | ***B. subtilis* OF5** | ***B. subtilis* OF4.2** |
| --- | --- | --- | --- |
| µ [h^-1^] | 0.24 ±0.02 | 0.17 ± 0.00 | 0.18 ± 0.02 |
| q_Glc_ [mmol g_CDW_^-1^ h^-1^] | 7.82 ± 0.77 | 6.78 ± 0.34 | 6.43 ± 0.67 |
| Y_X/Glc_ [g_CDW_ mol^-1^] | 30.95 ± 1.40 | 25.08 ± 1.18 | 28.03 ± 1.18 |
| Y_PGA/Glc_ [mol mol^-1^] | 0.38 ± 0.01 | 0.50 ± 0.00 | 0.34 ± 0.01 |
| Y_ACO/Glc_ [mol mol^-1^] | 0.02 ± 0.00 | 0.01 ± 0.00 | 0.03 ± 0.00 |
| Y_ACE/Glc_ [mol mol^-1^] | 0.18 ±0.01 | 0.03 ± 0.00 | 0.32 ± 0.02 |
| Y_LAC/Glc_ [mol mol^-1^] | 0.05 ±0.00 | 0.01 ±0.00 | 0.01 ± 0.00 |
| Y_BDO/Glc_ [mol mol^-1^] | 0.15 ±0.02 | 0.04 ±0.00 | 0.15 ± 0.02 |

**Table S2 – Metabolic reaction network used for flux estimation**

| **#** | **Genes** | **Reaction stoichiometry**  **(= reversible, > irreversible)** |
| --- | --- | --- |
| 1 | *ptsGHI* | GLC_EX + PEP > G6P + PYR |
| 2 | *pgi* | G6P = F6P |
| 3 | *pfkA* | F6P + ATP > FBP |
| 4 | *fbaA* | FBP = DHAP + G3P |
| 5 | *tpi* | DHAP = G3P |
| 6 | *gapA, gapB, pgk* | G3P = 3PG + NAD(P)H + ATP |
| 7 | *pgm, eno* | 3PG = PEP |
| 8 | *pyk* | PEP > PYR + ATP |
| 9 | *zwf, ykgB, gndA, yqeC, gntZ* | G6P > Ru5P + CO2 + 2 NADPH |
| 10 | *rpeA* | Ru5P = X5P |
| 11 | *ywlF* | Ru5P = Ri5P |
| 12 | *tktA* | Ri5P + X5P = S7P + G3P |
| 13 | *tal* | S7P + G3P = E4P + F6P |
| 14 | *tktA* | E4P + X5P = F6P + G3P |
| 15 | *pdhABCD, acoC, acoL, lpdV* | PYR > ACCOA + CO2 + NADH |
| 16 | *citA, citZ, citB* | ACCOA + OAA > ICIT |
| 17 | *icd* | ICIT > AKG + CO2 + NADPH |
| 18 | *odhAB, acoL, pdhD, lpdV, sucCD* | AKG > SUC + CO2 + ATP + NADH |
| 19 | *sdhABC* | SUC = FUM + QH2 |
| 20 | *fumC* | FUM = MAL |
| 21 | *mdh* | MAL = OAA + NADH |
| 22 | *mleA, ytsJ, malS, maeA* | MAL > PYR + CO2 + NAD(P)H |
| 23 | *pycA* | PYR + CO2 + ATP > OAA |
| 24 | *pckA* | OAA + ATP > PEP + CO2 |
| 25 | *ppc* | PEP + CO2 > OAA + Pi |
| 26 | *gltAB, glnA* | AKG + NADPH > GLU |
| 27 | *pgsBCA* | GLU + ATP > PGA |
| 28 | *ldh, lctP* | PYR + NADH > LAC |
| 29 | *alsS, ilvB, ilvH* | PYR + PYR > ACL + CO2 |
| 30 | *alsD* | ACL > ACO + CO2 |
| 31 | *bdhA* | ACO + NADH > BDO |
| 32 | *acoABCL* | ACO > 2 ACCOA + NADH |
| 33 | *pta, ackA* | ACCOA > ACE + ATP |
| 34 | *CO_2_ exchange* | CO2 = CO2_EX |
| 35 | *Biomass synthesis* | G6P_B + F6P_B + Ri5P_B + E4P_B + G3P_B + 3PG_B + PYR_B + ACCOA_B + AKG_B + OAA_B > BIOMASS |

**Table S3 – Anabolic demand.** Biomass precursor fluxes derived from growth-rate dependent biomass composition and strain-specific biomass yields.

| **Precursor** | **Anabolic demand [mol per 100 mol glucose]** | | |
| --- | --- | --- | --- |
|  | ***B. subtilis* PG10** | ***B. subtilis* OF4.2** | ***B. subtilis* OF5** |
| G6P | 1.88 | 1.83 | 1.66 |
| F6P | 0.01 | 0.01 | 0.01 |
| R5P | 1.62 | 1.36 | 1.20 |
| E4P | 1.31 | 1.15 | 1.02 |
| G3P | 1.10 | 1.10 | 0.99 |
| 3PG | 4.17 | 3.63 | 3.22 |
| PEP | 2.90 | 2.58 | 2.30 |
| PYR | 9.46 | 8.47 | 7.55 |
| ACA | 5.80 | 5.53 | 4.98 |
| 2-OX | 3.92 | 3.50 | 3.12 |
| OAA | 5.79 | 5.13 | 4.56 |

**Table S4 – Goodness of fit.** Experimentally determined and simulated relative fractions of mass isotopomers (MIDs) using an assumed measurement error of 0.3 mol%.

| **Fragment** | | ***B. subtilis* PG10** | | ***B. subtilis* OF4.2** | | ***B. subtilis* OF5** | |
| --- | --- | --- | --- | --- | --- | --- | --- |
|  |  | **exp.** | **sim.** | **exp.** | **sim.** | **exp.** | **sim.** |
| Ala 260 | m | 0.463 | 0.472 | 0.522 | 0.528 | 0.520 | 0.523 |
|  | m+1 | 0.393 | 0.387 | 0.347 | 0.341 | 0.346 | 0.341 |
|  | m+2 | 0.110 | 0.108 | 0.102 | 0.102 | 0.103 | 0.107 |
|  | m+3 | 0.034 | 0.033 | 0.029 | 0.029 | 0.031 | 0.030 |
| Ala 232 | m | 0.486 | 0.498 | 0.547 | 0.558 | 0.547 | 0.552 |
|  | m+1 | 0.406 | 0.397 | 0.353 | 0.346 | 0.353 | 0.350 |
|  | m+2 | 0.109 | 0.105 | 0.099 | 0.096 | 0.100 | 0.098 |
| Gly 246 | m | 0.747 | 0.750 | 0.747 | 0.747 | 0.738 | 0.734 |
|  | m+1 | 0.180 | 0.178 | 0.180 | 0.180 | 0.187 | 0.191 |
|  | m+2 | 0.073 | 0.072 | 0.073 | 0.073 | 0.075 | 0.075 |
| Gly 218 | m | 0.820 | 0.822 | 0.822 | 0.823 | 0.809 | 0.815 |
|  | m+1 | 0.180 | 0.178 | 0.178 | 0.177 | 0.191 | 0.185 |
| Val 288 | m | 0.292 | 0.291 | 0.369 | 0.367 | 0.359 | 0.359 |
|  | m+1 | 0.410 | 0.414 | 0.392 | 0.394 | 0.392 | 0.393 |
|  | m+2 | 0.214 | 0.213 | 0.173 | 0.174 | 0.178 | 0.179 |
|  | m | 0.063 | 0.063 | 0.051 | 0.051 | 0.054 | 0.054 |
|  | m+3 | 0.017 | 0.016 | 0.012 | 0.012 | 0.014 | 0.013 |
|  | m+4 | 0.004 | 0.003 | 0.003 | 0.002 | 0.004 | 0.002 |
| Val 260 | m | 0.294 | 0.299 | 0.371 | 0.379 | 0.360 | 0.370 |
|  | m+1 | 0.412 | 0.418 | 0.393 | 0.396 | 0.391 | 0.398 |
|  | m+2 | 0.214 | 0.209 | 0.172 | 0.167 | 0.176 | 0.172 |
|  | m+3 | 0.063 | 0.059 | 0.051 | 0.047 | 0.058 | 0.049 |
|  | m+4 | 0.017 | 0.015 | 0.013 | 0.011 | 0.015 | 0.011 |
| Leu 274 | m | 0.197 | 0.193 | 0.275 | 0.274 | 0.264 | 0.264 |
|  | m+1 | 0.370 | 0.376 | 0.385 | 0.391 | 0.383 | 0.390 |
|  | m+2 | 0.282 | 0.283 | 0.231 | 0.230 | 0.238 | 0.236 |
|  | m+3 | 0.113 | 0.112 | 0.082 | 0.080 | 0.086 | 0.084 |
|  | m+4 | 0.031 | 0.030 | 0.021 | 0.021 | 0.023 | 0.022 |
|  | m+5 | 0.007 | 0.007 | 0.005 | 0.004 | 0.005 | 0.004 |
| Leu 200 | m | 0.215 | 0.214 | 0.301 | 0.304 | 0.289 | 0.293 |
|  | m+1 | 0.392 | 0.401 | 0.403 | 0.412 | 0.402 | 0.411 |
|  | m+2 | 0.278 | 0.277 | 0.219 | 0.214 | 0.227 | 0.222 |
|  | m+3 | 0.093 | 0.089 | 0.062 | 0.058 | 0.066 | 0.062 |
|  | m+4 | 0.018 | 0.017 | 0.012 | 0.010 | 0.013 | 0.011 |
|  | m+5 | 0.004 | 0.003 | 0.003 | 0.001 | 0.004 | 0.002 |
| Ile 274 | m | 0.249 | 0.250 | 0.304 | 0.298 | 0.304 | 0.304 |
|  | m+1 | 0.391 | 0.398 | 0.386 | 0.392 | 0.384 | 0.392 |
|  | m+2 | 0.244 | 0.242 | 0.213 | 0.216 | 0.212 | 0.212 |
|  | m+3 | 0.086 | 0.083 | 0.073 | 0.073 | 0.074 | 0.071 |
|  | m+4 | 0.024 | 0.022 | 0.019 | 0.018 | 0.020 | 0.018 |
|  | m+5 | 0.006 | 0.004 | 0.005 | 0.004 | 0.006 | 0.003 |
| Ile 200 | m | 0.277 | 0.278 | 0.337 | 0.330 | 0.339 | 0.338 |
|  | m+1 | 0.415 | 0.422 | 0.405 | 0.411 | 0.404 | 0.410 |
|  | m+2 | 0.229 | 0.228 | 0.194 | 0.197 | 0.191 | 0.193 |
|  | m+3 | 0.062 | 0.060 | 0.051 | 0.051 | 0.051 | 0.049 |
|  | m+4 | 0.013 | 0.011 | 0.010 | 0.009 | 0.011 | 0.009 |
|  | m+5 | 0.003 | 0.001 | 0.003 | 0.001 | 0.005 | 0.001 |
| Pro 258 | m | 0.299 | 0.294 | 0.388 | 0.380 | 0.362 | 0.359 |
|  | m+1 | 0.409 | 0.414 | 0.390 | 0.394 | 0.392 | 0.397 |
|  | m+2 | 0.211 | 0.213 | 0.164 | 0.167 | 0.178 | 0.179 |
|  | m+3 | 0.064 | 0.063 | 0.047 | 0.048 | 0.054 | 0.053 |
|  | m+4 | 0.017 | 0.016 | 0.011 | 0.011 | 0.014 | 0.012 |
| Ser 390 | m | 0.424 | 0.413 | 0.465 | 0.461 | 0.467 | 0.459 |
|  | m+1 | 0.376 | 0.381 | 0.348 | 0.349 | 0.346 | 0.348 |
|  | m+2 | 0.147 | 0.151 | 0.140 | 0.141 | 0.140 | 0.144 |
|  | m+3 | 0.053 | 0.055 | 0.047 | 0.048 | 0.048 | 0.049 |
| Ser 362 | m | 0.455 | 0.445 | 0.498 | 0.494 | 0.501 | 0.490 |
|  | m+1 | 0.395 | 0.402 | 0.362 | 0.365 | 0.359 | 0.367 |
|  | m+2 | 0.150 | 0.154 | 0.141 | 0.141 | 0.140 | 0.142 |
| Ser 288 | m | 0.494 | 0.481 | 0.541 | 0.536 | 0.543 | 0.532 |
|  | m+1 | 0.390 | 0.399 | 0.351 | 0.356 | 0.349 | 0.360 |
|  | m+2 | 0.116 | 0.119 | 0.108 | 0.108 | 0.108 | 0.108 |
| Thr 404 | m | 0.313 | 0.318 | 0.347 | 0.351 | 0.340 | 0.344 |
|  | m+1 | 0.385 | 0.386 | 0.374 | 0.372 | 0.373 | 0.371 |
|  | m+2 | 0.201 | 0.199 | 0.190 | 0.188 | 0.192 | 0.192 |
|  | m+3 | 0.077 | 0.075 | 0.070 | 0.069 | 0.073 | 0.072 |
|  | m+4 | 0.023 | 0.022 | 0.020 | 0.020 | 0.022 | 0.021 |
| Thr 376 | m | 0.352 | 0.355 | 0.372 | 0.375 | 0.383 | 0.387 |
|  | m+1 | 0.392 | 0.392 | 0.377 | 0.378 | 0.373 | 0.373 |
|  | m+2 | 0.186 | 0.185 | 0.183 | 0.182 | 0.178 | 0.177 |
|  | m+3 | 0.070 | 0.068 | 0.067 | 0.065 | 0.066 | 0.063 |
| Phe 336 | m | 0.241 | 0.246 | 0.299 | 0.303 | 0.282 | 0.291 |
|  | m+1 | 0.378 | 0.393 | 0.376 | 0.384 | 0.365 | 0.380 |
|  | m+2 | 0.235 | 0.244 | 0.207 | 0.212 | 0.209 | 0.219 |
|  | m+3 | 0.086 | 0.088 | 0.074 | 0.076 | 0.078 | 0.082 |
|  | m+4 | 0.025 | 0.024 | 0.021 | 0.021 | 0.024 | 0.023 |
|  | m+5 | 0.017 | 0.005 | 0.012 | 0.004 | 0.019 | 0.005 |
|  | m+6 | 0.006 | 0.001 | 0.004 | 0.001 | 0.008 | 0.001 |
|  | m+7 | 0.006 | 0.000 | 0.004 | 0.000 | 0.007 | 0.000 |
|  | m+8 | 0.003 | 0.000 | 0.002 | 0.000 | 0.004 | 0.000 |
|  | m+9 | 0.002 | 0.000 | 0.002 | 0.000 | 0.003 | 0.000 |
| Phe 234 | m | 0.281 | 0.280 | 0.344 | 0.345 | 0.330 | 0.330 |
|  | m+1 | 0.408 | 0.420 | 0.396 | 0.405 | 0.392 | 0.404 |
|  | m+2 | 0.223 | 0.226 | 0.185 | 0.188 | 0.193 | 0.197 |
|  | m+3 | 0.063 | 0.061 | 0.052 | 0.050 | 0.056 | 0.056 |
|  | m+4 | 0.014 | 0.012 | 0.012 | 0.010 | 0.014 | 0.011 |
|  | m+5 | 0.004 | 0.002 | 0.004 | 0.001 | 0.005 | 0.002 |
|  | m+6 | 0.002 | 0.000 | 0.003 | 0.000 | 0.004 | 0.000 |
|  | m+7 | 0.002 | 0.000 | 0.003 | 0.000 | 0.003 | 0.000 |
|  | m+8 | 0.002 | 0.000 | 0.002 | 0.000 | 0.003 | 0.000 |
| Phe 302 | m | 0.728 | 0.720 | 0.727 | 0.723 | 0.724 | 0.721 |
|  | m+1 | 0.196 | 0.202 | 0.197 | 0.200 | 0.199 | 0.202 |
|  | m+2 | 0.076 | 0.078 | 0.076 | 0.077 | 0.077 | 0.077 |
| Asp 418 | m | 0.314 | 0.318 | 0.345 | 0.350 | 0.341 | 0.343 |
|  | m+1 | 0.383 | 0.385 | 0.373 | 0.371 | 0.370 | 0.371 |
|  | m+2 | 0.202 | 0.200 | 0.191 | 0.189 | 0.193 | 0.192 |
|  | m+3 | 0.077 | 0.076 | 0.071 | 0.070 | 0.074 | 0.072 |
|  | m+4 | 0.023 | 0.022 | 0.021 | 0.020 | 0.023 | 0.021 |
| Asp 390 | m | 0.353 | 0.354 | 0.374 | 0.374 | 0.385 | 0.386 |
|  | m+1 | 0.391 | 0.392 | 0.377 | 0.377 | 0.371 | 0.373 |
|  | m+2 | 0.187 | 0.186 | 0.183 | 0.183 | 0.178 | 0.177 |
|  | m+3 | 0.070 | 0.069 | 0.066 | 0.066 | 0.065 | 0.064 |
| Asp 316 | m | 0.386 | 0.389 | 0.408 | 0.411 | 0.422 | 0.425 |
|  | m+1 | 0.399 | 0.402 | 0.381 | 0.385 | 0.375 | 0.379 |
|  | m+2 | 0.161 | 0.160 | 0.158 | 0.158 | 0.152 | 0.152 |
|  | m+3 | 0.054 | 0.048 | 0.052 | 0.046 | 0.051 | 0.045 |
| Glu 432 | m | 0.214 | 0.216 | 0.255 | 0.256 | 0.261 | 0.261 |
|  | m+1 | 0.363 | 0.366 | 0.363 | 0.366 | 0.364 | 0.367 |
|  | m+2 | 0.260 | 0.258 | 0.238 | 0.236 | 0.235 | 0.233 |
|  | m+3 | 0.114 | 0.112 | 0.102 | 0.100 | 0.099 | 0.098 |
|  | m+4 | 0.039 | 0.038 | 0.033 | 0.033 | 0.033 | 0.032 |
|  | m+5 | 0.010 | 0.010 | 0.009 | 0.008 | 0.009 | 0.008 |
| Glu 330 | m | 0.275 | 0.281 | 0.361 | 0.363 | 0.342 | 0.343 |
|  | m+1 | 0.405 | 0.408 | 0.393 | 0.393 | 0.392 | 0.395 |
|  | m+2 | 0.227 | 0.222 | 0.179 | 0.178 | 0.190 | 0.189 |
|  | m+3 | 0.073 | 0.070 | 0.054 | 0.054 | 0.060 | 0.059 |
|  | m+4 | 0.020 | 0.018 | 0.013 | 0.013 | 0.016 | 0.015 |
| Lys 431 | m | 0.211 | 0.203 | 0.253 | 0.246 | 0.255 | 0.245 |
|  | m+1 | 0.355 | 0.360 | 0.360 | 0.363 | 0.358 | 0.361 |
|  | m+2 | 0.263 | 0.265 | 0.240 | 0.241 | 0.238 | 0.242 |
|  | m+3 | 0.118 | 0.118 | 0.103 | 0.104 | 0.104 | 0.106 |
|  | m+4 | 0.040 | 0.040 | 0.034 | 0.034 | 0.034 | 0.035 |
|  | m+5 | 0.011 | 0.011 | 0.009 | 0.009 | 0.009 | 0.009 |
|  | m+6 | 0.002 | 0.002 | 0.002 | 0.002 | 0.002 | 0.002 |
| Lys 329 | m | 0.237 | 0.239 | 0.278 | 0.284 | 0.289 | 0.291 |
|  | m+1 | 0.381 | 0.392 | 0.377 | 0.388 | 0.378 | 0.388 |
|  | m+2 | 0.252 | 0.249 | 0.230 | 0.224 | 0.223 | 0.220 |
|  | m+3 | 0.095 | 0.090 | 0.084 | 0.079 | 0.082 | 0.077 |
|  | m+4 | 0.028 | 0.025 | 0.024 | 0.021 | 0.023 | 0.020 |
|  | m+5 | 0.007 | 0.005 | 0.006 | 0.004 | 0.005 | 0.004 |
| Arg 442 | m | 0.178 | 0.179 | 0.199 | 0.196 | 0.205 | 0.208 |
|  | m+1 | 0.330 | 0.341 | 0.333 | 0.340 | 0.334 | 0.346 |
|  | m+2 | 0.269 | 0.276 | 0.261 | 0.267 | 0.256 | 0.261 |
|  | m+3 | 0.139 | 0.136 | 0.132 | 0.132 | 0.127 | 0.125 |
|  | m+4 | 0.055 | 0.050 | 0.052 | 0.048 | 0.051 | 0.045 |
|  | m+5 | 0.023 | 0.014 | 0.019 | 0.014 | 0.021 | 0.013 |
|  | m+6 | 0.007 | 0.003 | 0.005 | 0.003 | 0.006 | 0.003 |
| His 440 | m | 0.344 | 0.337 | 0.354 | 0.354 | 0.326 | 0.329 |
|  | m+1 | 0.360 | 0.372 | 0.351 | 0.356 | 0.353 | 0.355 |
|  | m+2 | 0.181 | 0.188 | 0.179 | 0.188 | 0.189 | 0.200 |
|  | m+3 | 0.077 | 0.075 | 0.078 | 0.074 | 0.085 | 0.082 |
|  | m+4 | 0.026 | 0.022 | 0.026 | 0.022 | 0.030 | 0.025 |
|  | m+5 | 0.009 | 0.005 | 0.009 | 0.005 | 0.012 | 0.006 |
|  | m+6 | 0.003 | 0.001 | 0.003 | 0.001 | 0.005 | 0.001 |
| Tyr 466 | m | 0.216 | 0.212 | 0.262 | 0.261 | 0.251 | 0.251 |
|  | m+1 | 0.361 | 0.363 | 0.361 | 0.361 | 0.354 | 0.357 |
|  | m+2 | 0.257 | 0.258 | 0.230 | 0.232 | 0.233 | 0.237 |
|  | m+3 | 0.112 | 0.114 | 0.098 | 0.101 | 0.102 | 0.106 |
|  | m+4 | 0.038 | 0.039 | 0.033 | 0.034 | 0.035 | 0.037 |
|  | m+5 | 0.010 | 0.010 | 0.008 | 0.009 | 0.010 | 0.010 |
|  | m+6 | 0.003 | 0.002 | 0.005 | 0.002 | 0.007 | 0.002 |
|  | m+7 | 0.001 | 0.000 | 0.002 | 0.000 | 0.003 | 0.000 |
|  | m+8 | 0.001 | 0.000 | 0.001 | 0.000 | 0.002 | 0.000 |
|  | m+9 | 0.000 | 0.000 | 0.000 | 0.000 | 0.001 | 0.000 |
| Tyr 302 | m | 0.726 | 0.720 | 0.724 | 0.723 | 0.721 | 0.721 |
|  | m+1 | 0.200 | 0.202 | 0.202 | 0.200 | 0.202 | 0.202 |
|  | m+2 | 0.074 | 0.078 | 0.074 | 0.077 | 0.076 | 0.077 |
| Rib 467 | m | 0.486 | 0.494 | 0.461 | 0.466 | 0.431 | 0.436 |
|  | m+1 | 0.291 | 0.280 | 0.299 | 0.293 | 0.312 | 0.302 |
|  | m+2 | 0.151 | 0.154 | 0.162 | 0.161 | 0.170 | 0.172 |
|  | m+3 | 0.053 | 0.053 | 0.057 | 0.058 | 0.060 | 0.064 |
|  | m+4 | 0.016 | 0.016 | 0.017 | 0.018 | 0.020 | 0.021 |
|  | m+5 | 0.004 | 0.004 | 0.004 | 0.004 | 0.006 | 0.005 |
| Rib 307 | m | 0.587 | 0.607 | 0.581 | 0.598 | 0.571 | 0.585 |
|  | m+1 | 0.270 | 0.255 | 0.275 | 0.261 | 0.282 | 0.268 |
|  | m+2 | 0.112 | 0.110 | 0.113 | 0.112 | 0.114 | 0.115 |
|  | m+3 | 0.031 | 0.029 | 0.031 | 0.030 | 0.032 | 0.031 |
| Glc 554 | m | 0.040 | 0.039 | 0.086 | 0.084 | 0.111 | 0.101 |
|  | m+1 | 0.498 | 0.498 | 0.465 | 0.464 | 0.447 | 0.435 |
|  | m+2 | 0.249 | 0.249 | 0.245 | 0.246 | 0.245 | 0.251 |
|  | m+3 | 0.145 | 0.146 | 0.139 | 0.139 | 0.136 | 0.141 |
|  | m+4 | 0.047 | 0.048 | 0.048 | 0.048 | 0.040 | 0.051 |
|  | m+5 | 0.017 | 0.016 | 0.014 | 0.016 | 0.015 | 0.017 |
|  | m+6 | 0.004 | 0.004 | 0.003 | 0.004 | 0.005 | 0.004 |
| Glc 319 | m | 0.648 | 0.647 | 0.622 | 0.620 | 0.597 | 0.599 |
|  | m+1 | 0.221 | 0.220 | 0.237 | 0.236 | 0.256 | 0.249 |
|  | m+2 | 0.103 | 0.104 | 0.108 | 0.110 | 0.111 | 0.115 |
|  | m+3 | 0.023 | 0.024 | 0.027 | 0.027 | 0.029 | 0.030 |
|  | m+4 | 0.006 | 0.006 | 0.006 | 0.006 | 0.007 | 0.007 |
| GlcN 553 | m | 0.078 | 0.081 | 0.137 | 0.140 | 0.149 | 0.145 |
|  | m+1 | 0.470 | 0.467 | 0.424 | 0.421 | 0.415 | 0.394 |
|  | m+2 | 0.250 | 0.247 | 0.244 | 0.242 | 0.244 | 0.251 |
|  | m+3 | 0.139 | 0.139 | 0.131 | 0.131 | 0.128 | 0.136 |
|  | m+4 | 0.045 | 0.047 | 0.046 | 0.047 | 0.045 | 0.052 |
|  | m+5 | 0.014 | 0.015 | 0.015 | 0.015 | 0.014 | 0.017 |
|  | m+6 | 0.004 | 0.004 | 0.004 | 0.004 | 0.004 | 0.005 |
| GlcN 319 | m | 0.613 | 0.611 | 0.582 | 0.581 | 0.564 | 0.564 |
|  | m+1 | 0.246 | 0.244 | 0.266 | 0.261 | 0.276 | 0.271 |
|  | m+2 | 0.108 | 0.111 | 0.114 | 0.118 | 0.118 | 0.122 |
|  | m+3 | 0.027 | 0.028 | 0.031 | 0.032 | 0.033 | 0.035 |
|  | m+4 | 0.006 | 0.007 | 0.007 | 0.008 | 0.007 | 0.008 |

**Table S5 – Relative and absolute net metabolic fluxes (in % and mmol g_CDW_^-1^ h^-1^)**

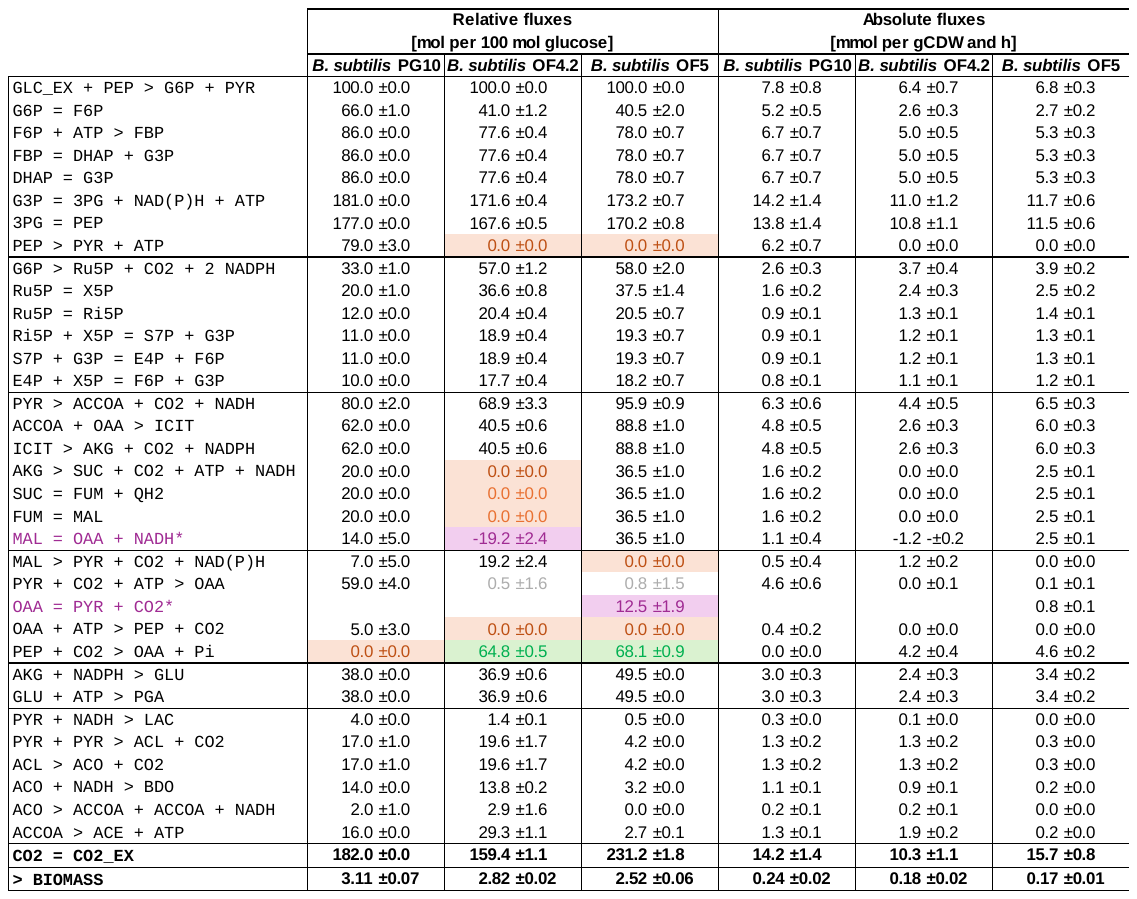
